# Insular cortex predictions regulate glucose homeostasis

**DOI:** 10.64898/2025.12.09.693191

**Authors:** Einav Litvak, Zhe Zhao, Inbar Perets, Ayal Lavi, Omer Izhaki, Rachel Barkan-Michaeli, Yishai Levin, Sarah A. Stern, Kfir Sharabi, Yoav Livneh

**Affiliations:** Department of Brain Sciences, Weizmann Institute of Science, Rehovot, Israel; Max Planck Florida Institute for Neuroscience, Jupiter, FL 33458, USA; Institute of Biochemistry, Food Science and Nutrition, The Robert H. Smith Faculty of Agriculture, Food and Environment, The Hebrew University of Jerusalem, Israel; de Botton Institute for Protein Profiling, The Nancy and Stephen Grand Israel National Center for Personalized Medicine, Weizmann Institute of Science, Rehovot, Israel

## Abstract

Brain-body interactions are essential for physical and emotional homeostasis. The brain uses information from the external world to predict upcoming bodily changes. This process involves interoceptive predictions, which are thought to play a central role in brain-body interactions. Yet there is little direct experimental evidence causally linking interoceptive predictions to regulation of bodily physiology. Here we address this by focusing on insular cortex and glucose homeostasis. We find that just before the onset of a meal, insular cortex exhibits a transient burst of activity, reflecting a prediction of the future metabolic state. This transient predictive burst of activity is essential for anticipatory insulin release, subsequent post-meal insulin release, post-meal glucose and lipid homeostasis, and post-meal metabolism signaling in the liver. Our results highlight that insular cortex predictive computations are essential for anticipatory physiological control and for subsequently maintaining metabolic homeostasis.

## Introduction

The brain and the body are in continuous dialog. Our brain constantly receives sensory information from within our body, as well as from the external environment, and then uses it to regulate bodily function^1–4^. Brain-body communication is essential for our physical and mental health, yet how it is achieved at the neurobiological level is still incompletely understood. Current models arising from fields of active inference and predictive processing propose that the brain computes interoceptive predictions about upcoming bodily changes and uses them to guide behavior and regulate bodily physiology^5–8^. However, there is very little direct experimental evidence causally linking active interoceptive inference to regulation of bodily function.

The insular cortex (InsCtx) encompasses within its mid-posterior division the primary visceral and gustatory cortex^9–11^. InsCtx receives internal sensory information regarding changes in bodily physiology, including energy and hydration state^12,13^, as well as external sensory cues^14–17^. Specifically in the context of feeding, InsCtx represents multimodal food-predicting cues, including visual, olfactory, and gustatory food related information^14–22^. Importantly, InsCtx activity is not necessary for normal taste function nor for feeding per se^15,18,23^. In contrast, InsCtx activity is necessary for learned food-cue guided behaviors, taste-related decision making^15,18,24,25^, and learned food aversion^26–30^. InsCtx has been suggested to compute predictive signals regarding upcoming physiological changes (i.e., interoceptive predictions^5,31,32^), but direct experimental evidence has been scarce. Moreover, we still lack direct evidence for the importance of such predictive signals for regulating bodily physiology. Interestingly, InsCtx is connected to different peripheral organs, as demonstrated by multi-synaptic pseudorabies tracing^33^.

Here, we set out to test the role of InsCtx predictions on bodily physiology. We reasoned that these predictions could drive anticipatory changes, including hormonal release, that bridge the long time-gap between the initial cortical prediction and subsequent metabolic changes, tens of minutes later. Indeed, a prevailing theme in physiological regulation is the pre-emptive response to anticipated physiological challenges^4,34,35^. For example, the sight, smell or taste of food drive specific physiological changes, termed “cephalic phase of digestion”, long before actual food ingestion and digestion^36,37^. Similarly, the sight and taste of water is used for pre-emptive renal regulation of fluid balance^38^. We chose to initially focus on anticipatory insulin (often called “cephalic phase insulin release”) as it is a hallmark of anticipatory physiological regulation^36^. Anticipatory insulin release is triggered by external food-related sensory stimuli such as the sight and smell of food, but primarily by taste^39,40^. It has been suggested to be essential for post-meal glucose homeostasis^41,42^, yet there has been conflicting evidence for its importance^39^. We thus hypothesized that InsCtx predictions could have an important role in regulating bodily physiology and metabolism through anticipatory physiological regulation, including anticipatory insulin release.

## Results

### InsCtx activity is necessary for learned anticipatory insulin release

Previous work suggested that anticipatory insulin release (also known as “cephalic phase insulin release”) can have a potentially “reflexive” component that is evoked by oral sugar sensing, as well as a learned component^39^. We reasoned that cortical predictions are more likely to be involved in the learned component. We therefore first sought to establish an assay for learned, non-reflexive, anticipatory insulin release. We note that we use here the term “anticipatory insulin release” to refer to the process often termed “cephalic phase insulin release”. That is, insulin release that is induced by food anticipation and the sensory properties of food, and precedes the post-meal rise in blood glucose levels^36,39,43^. We use the term “anticipatory” as we will relate it to anticipatory neural activity in InsCtx predicting the future post-meal metabolic state (see more below).

Anticipatory insulin release has been notoriously difficult to measure reliably^39^. We used within-subject measurements to overcome inter-subject variability. We tested various kinds of food and observed the most consistent and robust anticipatory insulin release with the highly caloric, sugar-containing milkshake, Ensure (**Supp. Fig. 1A**). We designed a simple protocol for freely behaving mice where we could separate the anticipatory (cephalic) phase of feeding from the consumption (meal) phase by placing Ensure in the homecage, allowing mice to approach it and then removing it upon first contact with it to measure anticipatory insulin release (i.e., first lick, similar to recent work that used chow^44^). We then returned the Ensure bowl to the homecage, allowed mice to consume it, and measured post-meal insulin levels (**Fig. 1A**). This method would thus induce sensory perception of food with visual, olfactory, and gustatory sensory stimuli, which together have been shown to evoke the strongest cephalic phase responses^37^. Based on previous work in mice^44,45^, we measured anticipatory insulin levels 1-3 minutes after removing the Ensure bowl to allow sufficient time for insulin levels to rise (see Methods). Our assay revealed that insulin release peaked at 1-3 minutes, declining thereafter (**Supp. Fig. 1B)**.

**Fig. 1:**
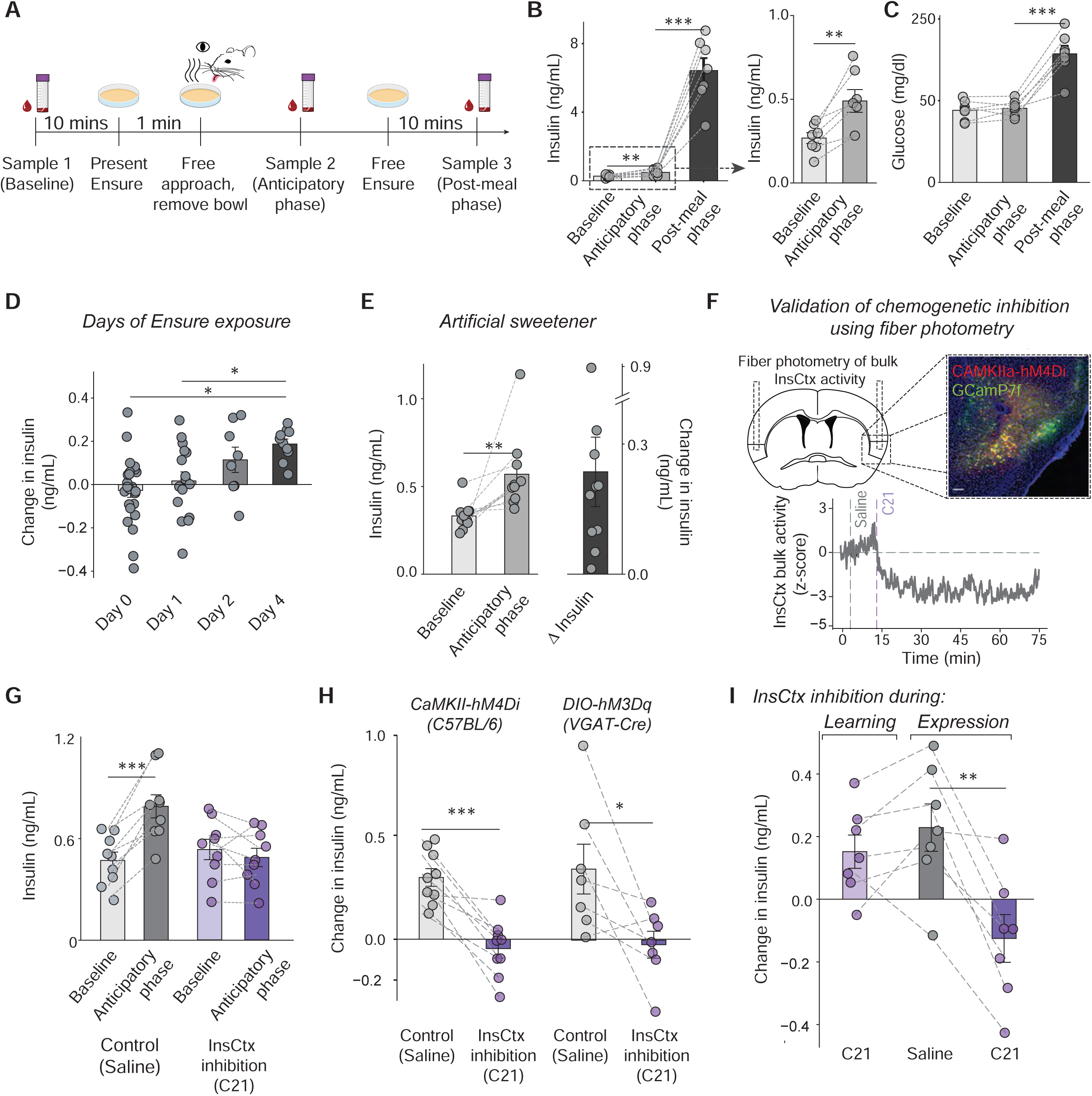
InsCtx activity is necessary for learned anticipatory insulin release. **(A)** Experimental design: Blood sampling protocol for measuring glucose and insulin at baseline (sample 1), anticipatory phase (sample 2) and post-meal (sample 3). Brown petri dish: bowl with 1 mL ensure. Mice approached it freely, and after they took one lick, we removed it. **(B)** *Left:* Insulin levels at baseline, during the anticipatory phase, and post-meal phase, after following the established protocol. *Right*: Zoom in to baseline and anticipatory phase insulin levels. There was a significant increase in anticipatory insulin (n=7, p = 0.002) and post-meal (p=0.00006). **(C)** Blood glucose values at baseline, during the anticipatory phase, and post-meal phase. There was no significant change during the anticipatory phase (n=7) and an expected increase after a meal (p=0.0001). **(D)** Change in anticipatory insulin levels (anticipatory phase - baseline) after different days of exposure to Ensure. Naïve mice (Day 0, n =24), and mice exposed for 1 day (n = 16), 2 days (n = 8), and 4 days (n = 10). We note that these experiments were performed with an across-subjects design, and experimental conditions therefore have different sample sizes (see Methods). Anticipatory insulin levels increased with repeated exposure. Mice with no previous exposure to Ensure had no anticipatory insulin response and 1 day of exposure was not sufficient to induce it (p = 0.80, day 1 vs day 0). Two days of exposure showed a trend towards an anticipatory insulin response (p = 0.14, day 2 vs day 0), with robust effects by Day 4 days compared to Day 0 (p = 0.002) and Day 1 (p = 0.03), and no statistically significant difference with day 2 (p=0.73). **(E)** Anticipatory insulin increases in the absence of sugar. Left and middle bars: anticipatory insulin significantly increased (n=10, p=0.001) compared to baseline when mice had experience with a milkshake-like drink containing sugar, but were exposed to the same drink with artificial sweetener instead of sugar on the test day. Also shown with the change in insulin (right bar). **(F)** Validation of chemogenetic inhibition efficacy and duration using fiber photometry recording of bulk InsCtx calcium signal. *Top left*: schematic of a coronal brain section at bregma, viral approach and fiber placement. *Top right*: example slice with co-expression of GCaMP7f and CaMKII-hM4Di-mCherry. Scale bar: 100 μm. *Bottom*: InsCtx bulk activity. Grey vertical line: intraperitoneal saline injection as a control, purple vertical line: intraperitoneal C21 injection. Note that InsCtx activity starts decreasing a few minutes after C21 injection and is maintained at low levels for at least an hour. **(G)** Chemogenetic inhibition of InsCtx attenuates anticipatory insulin release. Mice injected with AAV-CaMKII-hM4Di in the InsCtx had no increase in anticipatory insulin after injection of C21 compared to a saline injection (n=9, p_saline_ =0.00005). **(H)** Two different approaches to inhibit InsCtx show similar effects. Change in anticipatory insulin levels of the CaMKII-hM4Di cohort (inhibition of excitatory neurons, from ‘**G**’) and from hM3Dq in VGAT-Cre mice (activation of inhibitory interneurons), show that C21 injection attenuates anticipatory insulin release in response to Ensure presentation as compared to saline injection. (CaMKII-hM4Di: n=9, p=0.0002; VGAT-hM3Dq: n=7, p=0.02). **(I)** InsCtx activity is necessary for expression of learned Ensure associations for anticipatory insulin release, but not for learning them. *Left bar:* Mice received Ensure while InsCtx was chemogenetically inhibited (CaKMII-hMD4i) and still exhibited significant anticipatory insulin release upon presentation of Ensure (n=7, p=0.22). *Middle and right bars:* In the same mice, chemogenetic inhibition of InsCtx just before Ensure presentation (after experience with Ensure) significantly attenuated anticipatory insulin release compared to control saline injection (p=0.003). Data are presented as individual measurements (points) and mean (bars) ± SEM. Gray dashed lines represent the same mouse. P values were calculated by Student’s paired two sample t-test (mean comparison, for **B**, **C**, **G , H, I**), Wilcoxon signed rank test for **E**. ANOVA and Tukey HSD post hoc test for **D**. *p <0.05, **p <0.01, ***p <0.001, ****p <0.0001.

We observed an average two-fold increase in circulating insulin levels during the anticipatory phase, while blood glucose levels remained unchanged (**Fig. 1B,C**). In agreement with previous work, anticipatory insulin levels were substantially lower than post-meal insulin levels (glucose-stimulated insulin secretion; **Fig. 1B**, *Left*), but significantly higher than baseline levels^39,46^ (**Fig. 1B**, *Right*). This anticipatory insulin release was more prominent in fasted animals than in fed ones (**Supp. Fig. 1C**). Notably, while circulating insulin levels are known to undergo circadian regulation^47^, we did not find a significant correlation in our protocol between time-of-day and baseline insulin levels (**Supp. Fig. 1D**; see Methods), as well as with anticipatory insulin release levels or change in insulin (**Supp. Fig. 1E,F**). However, there was a positive yet non-significant correlation between baseline insulin and time-of day. We note that our analysis included only a 5-hour range and so we cannot exclude the possibility of circadian regulation of anticipatory insulin release on longer timescales that could be revealed using different protocols (e.g., at time differences of 12 hours).

Having established an assay for anticipatory insulin release, we next tested whether our assay captured the learned and/or reflexive component. We calculated the change in insulin levels for easier comparison between experimental groups. We did not observe anticipatory insulin release in mice encountering Ensure for the first time, although they did approach and lick the Ensure, suggesting the lack of anticipatory insulin release was not due to neophobia (**Fig. 1D**). In contrast, once we familiarized the same mice with Ensure in their homecage for 4 days, they exhibited robust anticipatory insulin release (**Fig. 1D**). Interestingly, 1 day of Ensure exposure was not sufficient to induce anticipatory insulin release. Although 2 days of experience with Ensure yielded higher anticipatory insulin in most of the mice, 4 days of exposure provided the highest and most consistent levels (**Fig. 1D**, see Methods). Thus, the observed anticipatory insulin release requires previous experience with Ensure, likely enabling the formation of expectations regarding the consequences of its ingestion. As such, our experimental protocol differs from previous work that showed that cephalic phase insulin release can have a potentially “reflexive” component that is evoked by oral sugar sensing^39^. To further test this distinction, we familiarized mice with a milkshake-like solution containing sugar for several days, but then replaced the sugar with artificial sweetener on the test day. This experiment showed that even in the absence of sugar, mice displayed robust anticipatory insulin release (**Fig. 1E**). Moreover, “reflexive” oral sweet sensing (in the absence of sugar) is not sufficient to induce anticipatory insulin release in our experiments, as we did not observe it in mice consuming Ensure without at least 2 days or prior experience with Ensure. Thus, anticipatory insulin release in our assay is learned and cannot be explained by “reflexive” oral sugar/sweet sensing. We speculate that this anticipatory insulin release requires associating the novel salient sensory properties of Ensure (smell and taste) with its post-ingestive consequences (energy and nutrients). Such associations between novel tastes or smells and their post-ingestive consequences have been suggested to be stored in the InsCtx^26,48,49^.

To test whether InsCtx activity is necessary for learned anticipatory insulin release, we inhibited its activity using chemogenetics. We injected AAV-CaMKII-hM4Di into InsCtx and verified that viral expression was not in inhibitory interneurons; **Supp. Fig. 1H-J**). We recorded bulk calcium signals using fiber photometry in awake behaving mice to verify that chemogenetic inhibition caused robust and prolonged (>1 hr) inhibition of InsCtx activity (**Fig. 1F**). We injected Compound 21 (C21, the chemogenetic agonist) or saline (counterbalanced for order, see Methods) 10 min before the baseline blood sample. Chemogenetically inhibiting InsCtx strongly attenuated anticipatory insulin release (**Fig. 1G**; **Supp. Fig. 1G**). Analysis of the change in insulin levels revealed a near-complete elimination of anticipatory insulin release (Control: 0.30 ± 0.04, C21: −0.05 ± 0.05; **Fig. 1H**). We further validated the importance of InsCtx for anticipatory insulin release using an alternative chemogenetic method of activating inhibitory interneurons via Cre-dependent hM3Dq expression in VGAT-Cre mice, observing similar attenuation of anticipatory insulin release (Control: 0.35±0.12, C21: −0.02±0.07, **Fig. 1H, Supp. Fig. 1I-J**). We note that InsCtx inhibition did not alter Ensure intake as mice consumed the same amount of Ensure (1 mL) in both experimental conditions (**Supp. Fig. 1K**). We did not observe such effects in control mice not expressing the chemogenetic proteins (**Supp. Fig. 1L**), and including non-responders^50,51^ in the analysis did not change these findings (**Supp. Fig. 1G**, see Methods).

Previous work has shown that InsCtx is necessary for both learning and expression of aversive taste-malaise associations^28,52^. We thus tested its role in our assay for learned anticipatory insulin release. We again used CaMKII-hM4Di to chemogenetically inhibit InsCtx during daily exposure to Ensure for 4 days but not during the test day in which we measured anticipatory insulin release. We found that mice still displayed anticipatory insulin release (**Fig. 1I**, *Learning*). In contrast, in the same mice, once we inhibited InsCtx during the test day, anticipatory insulin release was significantly attenuated (**Fig. 1I**, *Expression*). These data suggest that InsCtx is important for expression, but not for learning associations for anticipatory insulin release. Having established that InsCtx activity is necessary for anticipatory insulin release, we next sought to examine its activity during the anticipatory phase.

### InsCtx activity patterns at meal onset anticipate the future metabolic state

InsCtx responds to learned food cues that predict small amounts of food reward, especially in trial-structured behavioral tasks (e.g., cued Ensure drops or food pellets^15,24^). Previous theoretical and computational work has suggested that these responses could reflect predictive signals^5,31,32^. We have previously provided initial evidence for this idea^12^, but also found that in some cases, predictive activity patterns in InsCtx are not observed during ongoing self-paced consumption bouts^53^. We thus sought to examine InsCtx activity in the context of the anticipatory phase at the onset of a large meal. To do so, we used our previously established method of two-photon imaging of InsCtx activity via a microprism^15^. We imaged neuronal activity in layer 2/3 of mid-posterior InsCtx of head-fixed food-restricted mice just before and as they begin consuming Ensure ad libitum in the absence of any structured behavioral task (**Fig. 2A,B**). To create conditions that are similar to our protocol in freely moving mice, we initiated the protocol by delivering 1 drop of Ensure (∼5 µL), thereby providing initial olfactory cues, which then drive consumption. We used precise licking measurements to follow InsCtx activity before Ensure consumption and as mice began consuming it. Importantly, we repeatedly trained mice to consume Ensure ad libitum and then verified that Ensure anticipation in this head-fixed context also elicited anticipatory insulin release (**Fig. 2C**).

**Fig. 2:**
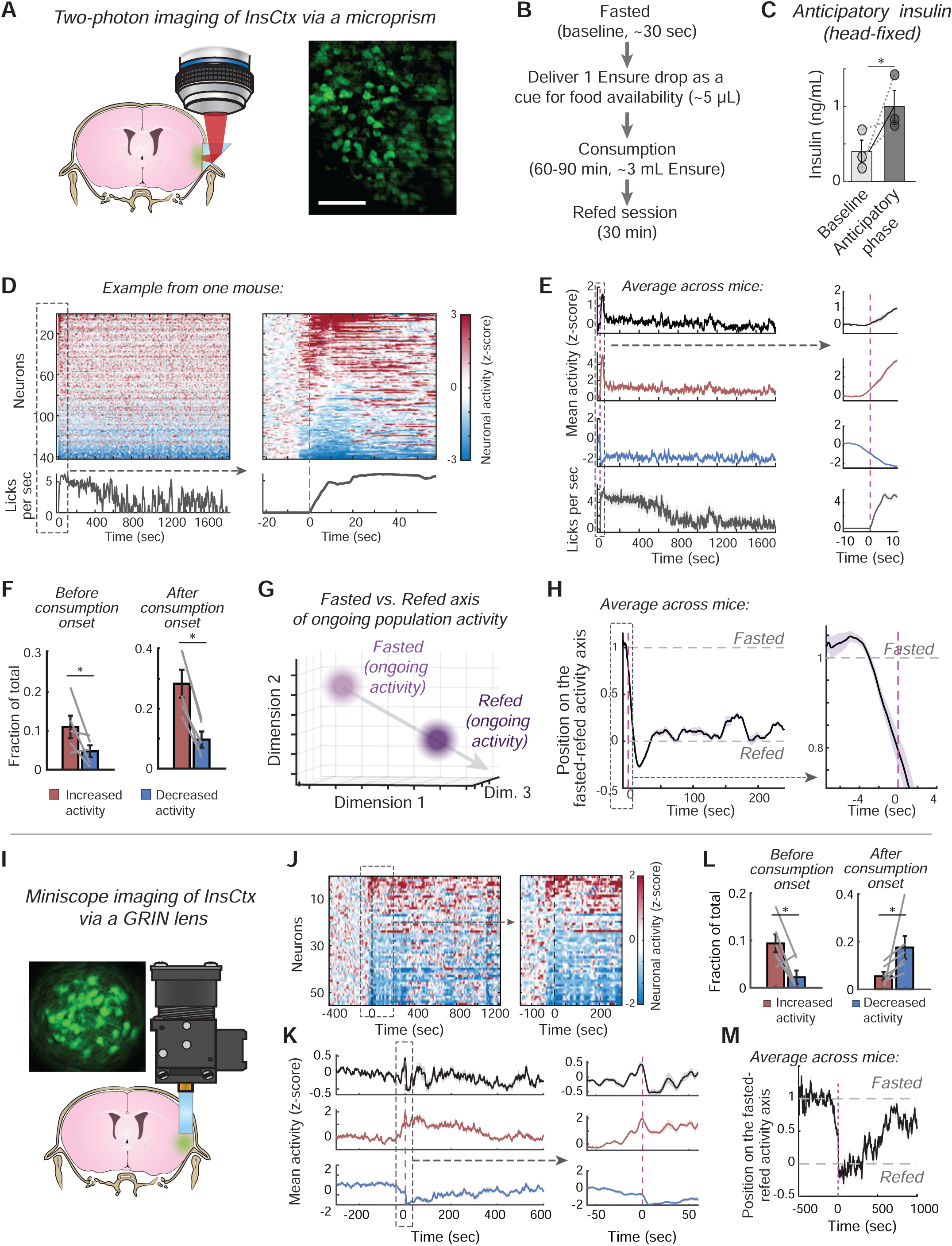
InsCtx activity patterns at meal onset anticipate the future metabolic state. **(A)** Two-photon calcium imaging of InsCtx through a microprism, schematic coronal brain section illustrating the approach, and example field-of-view. Scale bar: 100 µm. **(B)** Timeline of the head-fixed Ensure consumption two-photon imaging experiment. **(C)** Validation of anticipatory insulin release in head-fixed mice in anticipation of Ensure (p = 0.04). **(D)** *Top left:* example heatmap of all imaged InsCtx neurons from one mouse. *Bottom left:* licking behavior in the same experiment. *Right:* zoomed-in data from the dashed rectangle on the left side. **(E)** Same as ‘D’, but average across all mice. For each mouse we averaged either all neurons (black), or neurons with increased (red) or decreased (blue) activity. **(F)** Fraction of neurons with increased vs. decreased activity before (p=0.045) vs. at meal onset (p=0.001). **(G)** Schematic of the approach for assessing changes in the similarity of InsCtx activity patterns to ongoing activity patterns during fasted and refed states. **(H)** Pattern similarity of InsCtx population activity at meal onset to ongoing activity in the refed vs. fasted states. *Left*: first minutes. *Right*: first seconds. See Supp. Fig. 2 for the entire time-course. **(I)** One-photon miniscope calcium imaging of InsCtx through a GRIN lens, schematic coronal brain section illustrating the approach, and example field-of-view. **(J)** *Left:* example heatmap of all imaged InsCtx neurons from one mouse. *Right:* zoomed-in data from the dashed rectangle on the left side. **(K)** Same as ‘J’, but average across all mice. For each mouse we averaged either all responsive neurons, or neurons with increased or decreased activity. **(L)** Fraction of neurons with increased vs. decreased activity before (p=0.01) vs. at meal onset (p=0.04). **(M)** Pattern similarity of InsCtx population activity at meal onset to ongoing activity in the refed vs. fasted states. Values are Mean ± SEM across mice (two-photon imaging: n= 143±22 neurons per mouse from 5 mice; miniscope imaging: n= 51±20 neurons per mouse from 6 mice). P values were calculated by Student’s paired two sample t-test (mean comparison for **C**,**F**,**L**). *p<0.05, **p<0.01.

When examining the activity of InsCtx neuronal populations, we observed an initial burst of activity starting upon Ensure presentation approximately 5 sec before consumption onset and lasting ∼1 minute (**Fig. 2D**). This initial burst involved many InsCtx neurons increasing or decreasing their activity, with a slight bias to increased activity before consumption compared to baseline (**Fig. 2D-F; Supp. Fig. 2A**). Following the onset of consumption, activity became more biased to increased vs. decreased activity in terms of the number of neurons and their response magnitude (**Fig. 2F; Supp. Fig. 2A**). Therefore, although many neurons decreased and increased their activity before consumption onset compared to baseline, these changes were initially balanced at the population level such that the average activity across all neurons (i.e., with decreased and increased activity) appeared to mostly increase after consumption onset (**Fig. 2E**). This initial burst of InsCtx activity started before the onset of consumption, but did not reflect consumption or licking per se because: (1) it began before the first lick, and (2) it subsided after ∼1 min, when consumption was still ongoing and licking was still at peak rates (**Fig. 2D-E**).

We wondered whether the initial burst of activity represented a meaningful pattern, especially as it involved different neurons increasing or decreasing their activity, with a different balance between these responses across time. To test this, we used our previously established method to project neuronal population activity patterns onto a linear “fasted-refed axis”^12^ (**Fig. 2G**; see also ref. 54). We empirically defined this axis using activity patterns associated with a fasted state before meal onset (food restricted) and with a refed state after Ensure consumption^12^ (refed; **Fig. 2G,H**; **Supp. Fig. 2B**). Approximately 5 sec before consumption onset, activity patterns started to shift from a fasted-like pattern towards a refed-like pattern, reaching it within ∼30 sec (**Fig. 2H**). Interestingly, activity stabilized closer to a refed-like pattern after ∼1 min, even though mice were still licking and voraciously consuming Ensure, and average activity levels had returned to pre-meal baseline (**Fig. 2E,H**). In agreement with activity stabilizing after ∼1 min closer to a refed-like pattern, the change in activity before and at meal onset was correlated with activity 10 min later, albeit with much lower activity levels (**Supp. Fig. 2C**). Importantly, we performed control analyses in which we artificially increased InsCtx population activity, but this did not cause a similar shift towards refed-like patterns (**Supp. Fig. 2D**, see Methods). This suggests that the activity patterns we observed before and at meal onset are specific patterns that cannot be explained solely by overall increased activity.

Our anticipatory insulin experiments above were performed in freely-moving mice (**Fig. 1**). We thus tested whether the two-photon imaging results from head-fixed mice do indeed generalize to freely-moving mice. To test this, we implanted mice with mini-endoscopes in the mid-InsCtx and imaged 1-photon fluorescence during the same protocol for our anticipatory insulin measurements **(Fig. 2I**). Specifically, we recorded baseline activity for ∼5 min, then placed a bowl with Ensure in the cage and allowed mice to freely approach and consume it. Similar to the head-fixed two-photon imaging data, we found many InsCtx neurons either started increasing or decreasing their activity as mice were approaching the Ensure bowl to begin consuming Ensure, ∼20 sec before consumption onset (**Fig. 2J,K**). Average activity across all significantly responsive neurons increased as mice approached the Ensure bowl, and then returned to baseline after consumption began (**Fig. 2K**). This was due to more neurons being inhibited, but with stronger excitation in the excited neurons (**Fig. 2L; Supp. Fig. 2E**). We projected population activity on a “fasted-refed axis” in these data (**Fig. 2G**). Similar to the head-fixed two-photon data, activity patterns started to shift from a fasted-like pattern towards a refed-like pattern ∼20 sec before consumption began, as mice were approaching the Ensure (**Fig. 2M; Supp. Fig. 2F**). Activity stabilized in the refed-like pattern for ∼5 min and then gradually started shifting back towards the fasted-like pattern, even though average activity levels had returned to pre-meal baseline.

We wondered whether the differences between excited and inhibited neurons could be explained by differential activity of excitatory and inhibitory InsCtx neurons. To test this, we performed fiber photometry recording of bulk calcium dynamics in freely-moving mice from either inhibitory interneurons (in GAD2-Cre mice) or excitatory projection neurons targeted by retrograde AAV injections in InsCtx target regions^55^ (**Supp. Fig. 2G-H**). We found that the bulk activity of both populations increased as mice approached the Ensure bowl (**Supp. Fig. 2G-H**), similar to the average activity observed in the freely-moving cellular imaging data. Thus, the differences between neurons with increased or decreased activity cannot be explained by differential activity of excitatory and inhibitory InsCtx neurons, and is likely due to differential connectivity within the InsCtx microcircuit.

Together, these results show that during the anticipatory phase just before and at the onset of a meal, InsCtx activity patterns anticipate the future metabolic state. We therefore next tested to what extent these anticipatory activity patterns are involved in anticipatory insulin release.

### InsCtx anticipatory activity is necessary for anticipatory insulin release and for post-meal glucose homeostasis

To specifically test the importance of InsCtx anticipatory burst activity we used optogenetics for temporally precise inhibition. We used a similar protocol as above, but started optogenetic inhibition 15 seconds before presenting Ensure. We waited until mice took the first lick, immediately took away the bowl, and waited 3 minutes for insulin levels to rise^45^ (see Methods). After 3 minutes, we took the anticipatory phase blood sample and stopped illumination (**Fig. 3A**). We expressed ChrimsonR, a light-sensitive cation channel, or tdTomato (illumination control) in InsCtx inhibitory neurons using GAD2-Cre or VGAT-Cre mice (**Fig. 3B**), as previously established in other cortical areas and specifically also in InsCtx^56–58^. We validated that ChrimsonR expression in both mouse lines was predominantly in inhibitory interneurons (**Supp. Fig. 3A,B**). Importantly, we verified the efficacy and restricted duration of this optogenetic inhibition approach in awake behaving mice, finding that it drove time-locked overall inhibition of InsCtx activity. We did so by monitoring calcium dynamics specifically in excitatory projection neurons, targeted by retrograde AAV injections in InsCtx target regions^55^ (**Fig. 3C; Supp. Fig. 3C**). We verified that Ensure consumption, which took <5 min, was similar in all experimental conditions (**Supp. Fig. 3D**).

**Fig. 3:**
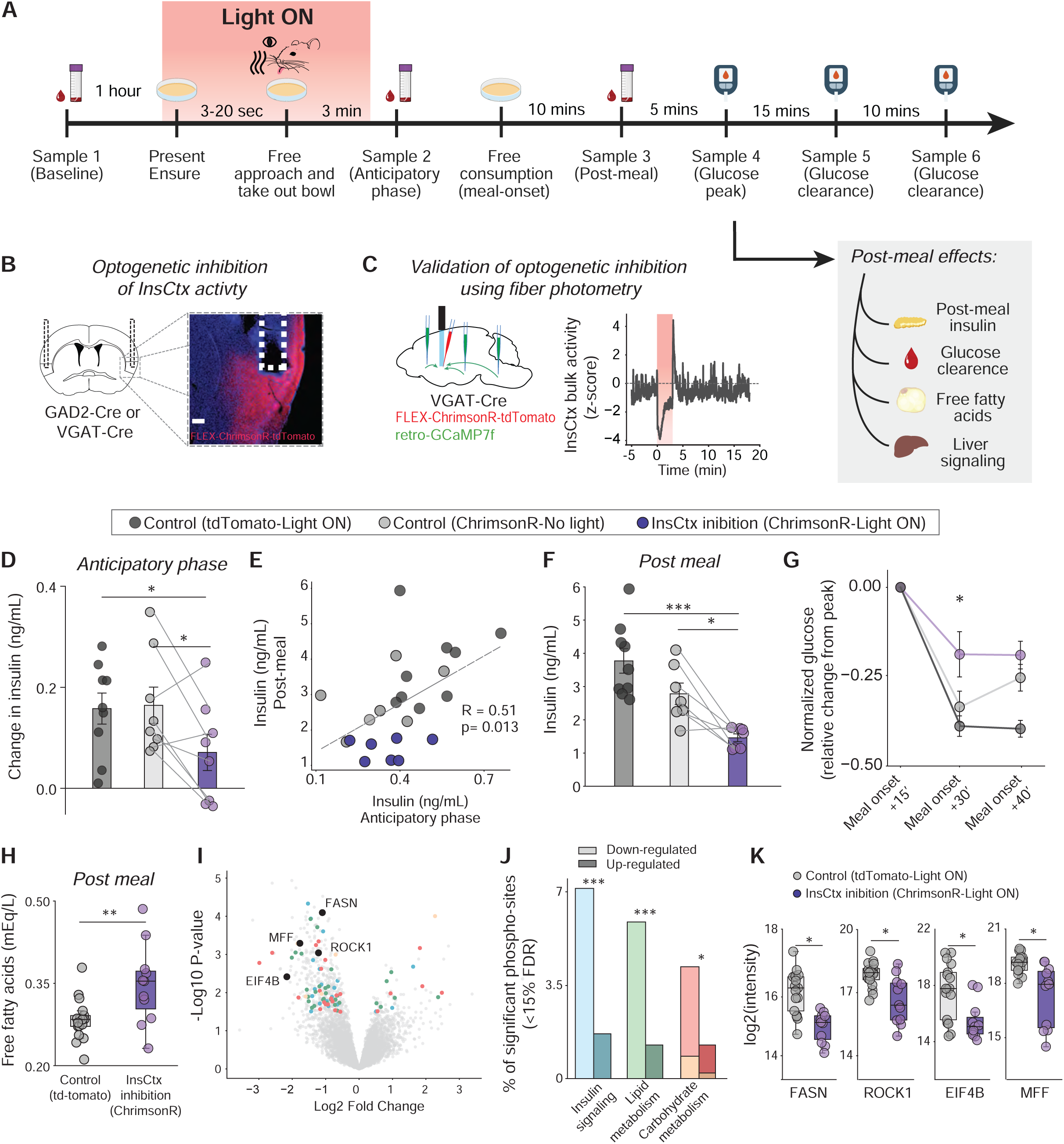
**InsCtx anticipatory activity is necessary for anticipatory insulin release and post-meal glucose homeostasis.** (**A**) Experimental design timeline. Blood sampling protocol for measuring glucose and insulin at baseline (sample 1), anticipatory phase (sample 2) and post-meal (sample 3). Brown petri dish: bowl with 1 mL Ensure. Fifteen seconds before Ensure presentation, we turned on the LED. Mice approached the Ensure freely, and after taking one lick, we removed it. We returned it after taking sample 2 and turned off the light. Then, 10 minutes later, we collected sample 3. In the within-subjects experiment, we collected three extra timepoints of blood glucose levels with a glucometer to build a glucose clearance curve. In the across-subject experiment, 15 minutes after returning the Ensure for free consumption, we collected liver tissue and whole blood (Light grey square). (**B**) *Left:* Schematic of a coronal brain section at bregma, area of injection and fiber placement. *Right:* Example section with InsCtx ChrimsonR expression and fiber placement. Scale bar: 200 μm. (**C**) Validation of optogenetic inhibition efficacy and duration using fiber photometry. *Left*: schematic of a sagittal brain section at bregma, viral approach with retro-GCaMP7f (green pipettes) injected in mid-InsCtx target regions (anterior InsCtx, central amygdala and basolateral amygdala, and rostral nucleus of the solitary tract), ChrimsonR (red pipette) injected into mid InsCtx, and fiber placement. *Right*: InsCtx bulk calcium activity, red square: optogenetic inhibition period of 3 minutes. Inhibition of InsCtx starts immediately after light is switched on, lasts for the whole illumination period and recovers to baseline after light is switched off. (**D**) Change in insulin demonstrating the extent of the effect of the inhibition compared to the illumination control (Light ON control n_chrimson_=8, n_tdTomato_=9 p=0.04) and the No light control (No light control n=8, p=0.02), with no difference between both control groups (p=0.92). **(E-F)** Reducing anticipatory insulin release affects post-meal insulin release. (**E**) The reduction of post-meal insulin is significantly correlated to the reduction in anticipatory insulin (n=16, R = 0.51, p = 0.013). (**F**) Post-meal insulin release is reduced after InsCtx anticipatory activity is inhibited compared to illumination control (tdTomato, Light ON, n=9, p=0.0003) and to No light control (ChrimsonR, No light, n=7, p = 0.02). No difference between both control days (p=0.07). We note that one mouse was excluded from this analysis due to post-meal blood sample degradation. (**G**) Change in glucose levels relative to the peak (15 minutes after meal onset). Glucose takes longer to be cleared from the blood after a meal when anticipatory insulin was reduced, compared to the illumination control (n_tdTomato_=9, p=0.02), and to the ‘no light’ control (Chrismon, No light, n_chrimson_=8, p = 0.02), with no difference between control groups (p=0.61). (**H**) Post-meal reduction in free fatty acids measured 15 minutes post meal onset (25 minutes after end of optogenetic inhibition). There is a lower post-meal reduction in free fatty acids compared to control, leading to significantly higher levels (n_tdTomato_ = 17, n_chrimson_ = 11, p = 0.009). **(I-K)** Post-meal liver phospho-proteomics analysis following InsCtx optogenetic inhibition during the anticipatory phase. (**I**) Volcano plot of up-regulated and down-regulated phosphosites at 15 minutes post meal onset. Overview of phosphosites of proteins related to select metabolism categories: Insulin signaling (blue), Lipid metabolism (green), Carbohydrate metabolism (red) and Gluconeogenesis (yellow). Significantly down-regulated phosphosites of relevant proteins related to these terms are highlighted in black (see ‘K’ for p-values). (**J**) Percentage of significant phosposites related to different aspects of metabolism out of all significant phosphosites (FDR<15%), same color references as in ‘G’, light shade indicates down-regulation and darker shade up-regulation. Phosphosites related to insulin signaling, lipid metabolism, and carbohydrate metabolism showed a significant bias towards down-regulated phosphorylation compared to a background distribution (Fischer’s exact-test. p_insulin_ = 0.0001, p_lipids_ = 0.00008, p_CH_ = 0.02). (**K**) Example of individual significantly down-regulated phosphosites of proteins related to metabolism (n_tdTomato_ = 17, n_chrimson_ = 11): FASN: adjusted p = 0.01, ROCK1: adjusted p = 0.02, EIF4B: adjusted p = 0.047, MFF: adjusted p = 0.02. FDR 5%=0.0541. Data are presented as individual measurements (points) and mean (bars or box plots) ± SEM. Gray dashed lines represent the same mouse. P values were calculated with Student’s paired two sample t-test (for **D**), Student’s independent two sample t-test (for **F,G,H,K**) Wilcoxon signed-rank (for **F,G**), Fischer’s exact-test (for **J**), Spearman’s rank test (for **E**). *p<0.05, ** p <0.01, ***p <0.001, ****p <0.0001.

Using this approach, we found that time-locked optogenetic inhibition of InsCtx only during initial Ensure presentation strongly attenuated anticipatory insulin release (Control tdTomato, Light ON: 0.16±0.03; Control ChrimsonR, No light: 0.16±0.04; InsCtx Inhibition: 0.07±0.04; **Fig. 3D**). The rapid, preparatory increase in circulating insulin levels prior to food consumption was not affected by InsCtx light illumination in control mice expressing tdTomato instead of ChrimsonR (**Fig. 3D**). The effects of optogenetic inhibition did not depend on the order of the experimental conditions (**Supp. Fig. 3E**, see Methods), and it was not affected by including non-responders^50,51^ in the analysis (**Supp. Fig. 3F**, see also **Supp. Fig. 3G,H** for the post-meal effect described below; see Methods).

We note that we had to modify our protocol to include longer waiting periods for habituation to the optic fiber patch-cord in our time-locked optogenetic approach. These longer waiting periods tended to elevate blood glucose levels in all experimental groups, which was likely due to associated stress, a well-known factor causing blood glucose level elevation^59^ (**Supp. Fig. 3I; Supp. Fig. 4A**; also compare to **Fig. 1C**). Indeed, we observed elevated blood glucose also in a separate group of mice that did not exhibit elevated anticipatory insulin (**Supp. Fig. 3J,K**, see Methods). Control analyses further confirmed that there was no significant correlation between glucose levels and anticipatory insulin in any of our chemogenetic and optogenetic assays and in all experimental groups (**Supp. Fig. 3L** and **Supp. Fig. 4B,C** for optogenetic experiments; **Supp. Fig. 1M** for chemogenetic experiments). Thus, the anticipatory increase in insulin we observed cannot be attributed to changes in blood glucose levels.

These data demonstrate that InsCtx anticipatory burst pattern activity is essential for anticipatory insulin release. Moreover, we establish a way to specifically inhibit the neural control of the anticipatory cephalic phase responses, including anticipatory insulin release. Due to the restricted duration of our optogenetic inhibition (**Fig. 3C**), inhibiting InsCtx only at meal onset likely does not affect subsequent InsCtx activity associated with consumption of the meal itself and with post-meal digestion. We therefore next tested whether inhibiting InsCtx only at meal onset would have consequences on post-meal glucose homeostasis (**Fig. 3A**).

We first examined the relationship between levels of anticipatory insulin and post-meal insulin, and found they were significantly correlated (**Fig. 3E; Supp. Fig. 3G**), suggesting that inhibiting InsCtx also affected post-meal insulin by attenuating anticipatory insulin levels. We tested this more directly by comparing post-meal insulin levels 10 min after meal onset, with and without InsCtx inhibition during the anticipatory phase, and indeed found it was significantly lower (**Fig. 3F; Supp. Fig. 3H**).

We reasoned that lower post-meal insulin levels could have consequences on post-meal circulating glucose levels^41,42^. Indeed, we found that inhibiting InsCtx only during the anticipatory phase affected glucose clearance. The within-subject normalized change in glucose (relative to the glucose peak at 15 min after meal onset) showed ∼50% less glucose clearance 30 min after meal onset, following InsCtx inhibition during the anticipatory phase (**Fig. 3G; Supp. Fig. 3M**). This was not due to differences in the amount of Ensure consumed (**Supp. Fig. 3D**). These results are consistent with the lower post-meal insulin levels (**Fig. 3F**). However, since glucose metabolism involves integration of multiple physiological changes, the reduced glucose clearance could additionally involve other physiological factors beyond insulin that were affected by InsCtx inhibition during the anticipatory phase 30 min earlier.

To further examine the consequences of inhibiting InsCtx in the anticipatory phase in acute experiments that require cross-subject comparisons (e.g., when collecting peripheral tissue), we repeated the experiment using two new different cohorts of mice: a control group (expressing tdTomato in InsCtx, n=17) and the experimental group (expressing ChrimsonR, n=11). Although the post-meal effects we examined might not necessarily be dependent on anticipatory insulin alone^60^, we verified that there was a decrease in anticipatory insulin only in the ChrimsonR group, but not in the tdTomato group (**Supp. Fig. 4D-F**). Ensure consumption was similar for both cohorts (**Supp. Fig. 4G**). Interestingly, in line with the observed decrease in post-meal insulin levels and post-meal glucose clearance, we also found significantly higher post-meal levels of circulating free fatty acids, suggesting reduced parasympathetic inhibition of adipose tissue lipolysis, and/or increased sympathetic output driving lipolysis^61^ (**Fig. 3H**).

In the same mice, we also examined post-meal liver signaling using phospho-proteomics (**Fig. 3I-K**). We chose to examine phosphorylation events due to the time-course of our experiment, which is potentially too rapid to induce large changes in gene expression. We first identified the optimal post-meal time-point for identifying relevant phosphorylation changes in the liver using western blot analysis (**Supp. Fig. 4H**). Then, using phospho-proteomic analysis we found substantially more downregulated vs. upregulated phosphorylation sites (85.1% vs. 14.9%, p < 4.75e-40 vs. background; **Fig. 3I**; **Supp. Fig. 4I)**, suggesting alteration of post-meal signaling. We examined the changes in phosphorylation among 4 relevant metabolism categories: insulin signaling, lipid metabolism, carbohydrate metabolism, and gluconeogenesis (**Fig. 3I,J**). We performed a permutation test to empirically assess the false discovery rate (FDR) and to set significance thresholds (5% FDR = 0.05 and 15% FDR = 0.13) (**Supp. Fig. 4J**). Within some metabolism categories, we found a bias to downregulated vs. upregulated phosphorylation sites: insulin signaling (7.13% vs. 1.68%, p < 0.0001 vs. background), lipid metabolism (5.87% vs. 1.26%, p < 0.00008 vs. background), and carbohydrate metabolism (4.19% vs. 1.26%, p < 0.02 vs. background), but not for a subset of carbohydrate metabolism terms related to gluconeogenesis (0.83% vs. 0.21% %, p < 0.2 vs. background; **Fig. 3J**; **Supp. Fig. 4K**). These data suggest that inhibiting InsCtx during the anticipatory phase had a stronger inhibitory effect on post-meal liver insulin signaling, lipid metabolism, and carbohydrate metabolism, but potentially not related to gluconeogenesis. This could indicate that the higher circulating glucose levels we observed might not be due to modified liver gluconeogenesis.

Upon closer examination of individual affected phosphorylation sites, we observed down-regulation of central regulators of lipid synthesis, such as fatty acid synthase (FASN) and rho-kinase 1 (ROCK1) (**Fig. 3K**). Consistent with previous work on anticipatory regulation of liver function likely independent of insulin action^60^, we also found down-regulation of protein translation machinery, such as the translation initiation factors eIF4b (**Fig. 3K**) and eIF3a (**Table S1**), as well as ribosomal protein S6 kinase A4 (RPS6KA4; **Table S1**). Furthermore, we observed a down-regulation in phosphorylation of serine 131 in mitochondrial fission factor (MFF), a central regulator of mitochondrial organization, important for systemic and hepatic glucose metabolism^62^ (MFF-S131; **Fig. 3K**). MFF-S131 has been recently shown to have an important role in anticipatory regulation of liver function to maintain glucose homeostasis^63^. These results suggest that InsCtx inhibition during the anticipatory phase could have also affected liver function and glucose homeostasis through other parallel pathways, independent of insulin.

In summary, inhibiting InsCtx activity only during the anticipatory phase resulted in several post-meal changes 10-30 min later (each measured at the relevant post-meal time-point). Together, these results show that InsCtx anticipatory activity at meal onset represents a potential prediction of future metabolic state. This putative cortical prediction is essential for anticipatory insulin release, and has an important contribution for post-meal glucose homeostasis.

## Discussion

The past years have seen substantial progress in our understanding of the neurobiology of brain-body communication, especially from the perspective of peripheral organ innervation, brainstem sub-populations, and hypothalamic sub-populations^64–71^. Yet one less explored aspect of brain-body communication has been the link between higher-order brain function, such as learning, memory, and predictive processing, to bodily physiology. Such higher-order processes likely rely on multiple interacting brain regions, with important hubs such as the InsCtx and anterior cingulate cortex^70^. Much less progress has been made in understanding the connection between neural computations related to higher-order brain function in these brain regions and bodily physiology.

Here, we begin to fill this gap by focusing on the InsCtx and anticipatory physiological regulation in the context of the cephalic phase of digestion. We initially focused on anticipatory hormonal release as a way to bridge the long time-gap between the pre-meal cortical prediction and post-meal bodily changes, which occur tens of minutes later. Nevertheless, other possible mechanisms could work in parallel to bridge this long time-gap. Previous work has suggested that cephalic phase insulin release has a reflexive component that is evoked by sweet taste, and specifically sensing of glucose on the tongue^39,72^. Interestingly, our behavioral model required prior experience with Ensure, suggesting that the sugar present in Ensure was not sufficient to engage the reflexive cephalic phase insulin release. This could be due to the very small amount of Ensure that mice consume in our assay (∼1 lick) as compared to previous taste-induced assays that involved consumption of much larger quantities^45^. Instead, our protocol required daily experience with Ensure for a few days, to associate its salient smell and taste with post-ingestive nutrient absorption and energy repletion. InsCtx activity was necessary for retrieval of these learned associations, but not for the learning process itself. We suggest that the cortical prediction adds an important layer on top of the reflexive component, thereby allowing more flexibility in a changing environment, as well as an earlier preparatory physiological responses.

Since InsCtx inhibition affected a wide range of post-meal changes, we propose that InsCtx predictions could be essential for many other aspects of anticipatory physiological regulation in the context of food anticipation, but also in other contexts such as fluid anticipation during dehydration^12,73^, and immune responses^74,75^. Indeed, our results are complementary to a recent study showing how food odor promotes systemic lipid utilization^76^. We thus speculate that InsCtx connections with the olfactory system could play a role in anticipatory regulation of lipid utilization.

InsCtx is also multi-synaptically connected to peripheral organs, via its projections to brainstem vagal motor neurons^77^. Multi-synaptic pseudorabies tracing suggests that these vagal motor neurons could include those that innervate the pancreas^33^. Future work could reveal how InsCtx may regulate different known sympathetic and parasympathetic pathways to modulate multiple aspects of bodily physiology^10,78^. This could also include under-explored InsCtx interactions with the enteric nervous system and pancreatic alpha cells, whose secretory functions are also an important aspect of glucose homeostasis^79^. Furthermore, future work could reveal how the InsCtx-to-brainstem pathway interacts with hypothalamic pathways that regulate similar physiological processes of energy and fluid balance^38,44,60,63,66–69,80,81^. Finally, future work could reveal whether different InsCtx activity patterns predict specific aspects of the metabolic state, or whether they predict a coarse-grained integration of many converging signals^82^. Linking cortical predictive computations to the above questions could have important translational implications due to the increasing worldwide prevalence of obesity, diabetes and other metabolic disorders.

## Acknowledgements

We thank members of the Livneh lab, Henning Fenselau, and Daphna Nachmani for invaluable discussions. We thank Michael Walker and Marc Donath for advice on optimizing anticipatory insulin release assays. This work was supported by research grants from the European Research Council (ERC StG #101039145; to YL), Israel Science Foundation (ISF #3015/22; to YL), and the Center for New Scientists of the Weizmann Institute of Science (to YL), NIH DP2DA060436 (to SAS) and the Max Planck Society (to SAS).

## Methods

All animal care and experimental procedures were approved by the Institutional Animal Care and Use Committee. Sample sizes were chosen to reliably measure experimental parameters, while remaining in compliance with ethical guidelines for minimizing animal use and keeping with standards in the relevant fields. Experiments did not involve experimenter-blinding. But post-hoc histological analysis of viral expression and optic fiber tract targeting we performed with blinding. In within-subject comparisons, animal subjects were not randomly allocated to experimental groups. In between subject comparisons, animals were randomly allocated to experimental and control groups.

### Surgical procedures

#### Stereotaxic injections

Mice were 8-23 weeks old at the time of injection. Mice were anesthetized with isoflurane in air (induction, 3%; maintenance, 1%–2%), and placed into a stereotaxic apparatus (RWD). After exposing the skull via a small incision, a small hole was drilled for injection. A pulled-glass pipette with 20–40 μm tip diameter was inserted into the brain, and virus was injected using an air pressure system (PV850, WPI). The pipette was withdrawn 3 min after injection. For postoperative care, mice were injected subcutaneously with slow-release buprenorphine (LA) (0.00325 mg/kg, VetMarket) and carprofen (5mg/kg, Dechra).

We used the following volumes of virus and injection coordinates: InsCtx (200 nl, Bregma: AP: +0.3mm and - 0.3mm for chemogenetics, and 0.0mm for optogenetics, DV: −4.0, ML: ∼4.0 mm), Central Amygdala (CeA; 200 nl, AP: −1.4mm, DV: −4.6 mm, ML: ±2.5 mm), Dorsal Vagal Complex (DVC; 200 nl, AP: −6.7mm, DV: −4.2 mm, ML: ±1.05 mm). All injections were done bilaterally. For optogenetic inhibition control experiment and fiber photometry of excitatory neurons, retroAAV-GCaMP7f: Anterior InsCtx: AP: +1.7mm, DV: −3.5 mm, ML: 3.20 mm, CeA and basolateral amygdala: AP: −1.4mm, DV: −4.6 to −4.4 mm, ML: 2.9 mm, DVC: AP: −6.7mm, DV: −4.5 mm, ML: 1.05 mm

We used the following viruses, all from Addgene: AAV8-hSyn-GCaMP7f, AAV1-hDlx-hM3Dq-nls-dTomato, AAV8-CaMKII-hM4Di–mCherry, AAV5-FLEX-ChrimsonR-tdTomato, retroAAV-hSyn-jGCaMP7f, AAV9-hDlx-FLEX-tdTomato, AAV8-hSyn-DIO-hM3D(Gq)-mCherry, AAV8-hSyn-FLEX-jGCaMP7f-WPRE, AAV8-CAG-tdTomato. We used AAV1-hSyn-GcaMP6f for two-photon imaging and AAV1-hSyn-GcaMP8m for miniscope imaging. Mice used for in vivo two-photon imaging were instrumented with a headpost and a 2 mm microprism, centered over the mid InsCtx. Implantation of microprisms (2 mm prisms; #MCPH-1.0; Tower Optical; coated with aluminum along their hypotenuse) were performed as previously described^15^. For GRIN lens the AAV was loaded in a glass pipette and infused into the brain by a nanoliter injector (DRUMMOND, nanoinject II). The coordinates were: AP: +0.5 mm, ML: 3.85 mm, DV: 3.85 mm (Inscopix, 0.5mm X 6.1mm, 1050-004415).

#### Chemogenetic experiments

C57BL/6 mice or VGAT-cre (JAX # 028862) were stereotaxically injected with AAV8-CaMKII-hM4Di–mCherry or AAV8-hSyn-DIO-hM3D(Gq)-mCherry, respectively.

#### Optic fiber implantation for fiber photometry and optogenetics

For validation of chemogenetic inhibition, we used chemegenetic virus and GCaMP7f. For validation of optogenetic inhibition, we used ChrimsonR and retro-GCaMP7f. We injected retro-GcaMP into several InsCtx target regions (see details above) in order to selectively express GcaMP in excitatory projections neurons across InsCtx layers.

First, C57BL/6 mice, GAD2-cre (JAX #01080), or VGAT-Cre (JAX #028862) mice (C57BL/6 background) were stereotaxically injected with the relevant virus as specified above. Then, an optic fiber (400-μm diameter core; Multimode; NA 0.50; 5mm length, Thor Labs or RDW) was implanted over the InsCtx (same coordinates as injection except D/V = 3.95). The fiber was fixed to the skull using C&B Metabond (Parkell). Mice were allowed at least 2 weeks for recovery before habituation started.

### Fiber photometry

To reliably assess activity levels while minimizing potential motion artifacts, in vivo assessments of chemogenetic or optogenetic InsCtx inhibition were performed in head-fixed mice. Mice were habituated to head-fixation 2 days before recording InsCtx bulk calcium signal by placing them in the rig, connecting the optogenetic patch-cords (in the case of the optogenetic inhibition control) for increasing amounts of time between 10, 20, 60 or 75 minutes for the chemogenetic control or 10, 20 or 40 minutes for the optogenetic control.

On the day of fiber photometry recording, mice were placed on the rig and connected to the patch-cords. We recorded the fiber photometry signal from the left hemisphere while inhibiting both hemispheres. For chemogenetics, we recorded 5 minutes of baseline activity, followed by intraperitoneal saline injection. We recorded for 10 more minutes and then injected C21 intraperitoneally before recording for an additional 60 minutes. For optogenetics, we recorded 5 minutes of baseline activity, and then turned on the LED light (∼625 nm, 10 mW to each hemisphere; 10Hz with 20% duty cycle; Orange-red LED, prizmatix) for 3 minutes of recording, turned it off and recorded for 15 more minutes.

For measuring activity levels during approach to Ensure in freely-moving mice, we used two cohorts: C57BL/6 mice injected with retroAAV-GCaMP6f in InsCtx projection targets to retrogradely label excitatory neurons, or GAD2-Cre mice injected with FLEX-GCaMP7f to target inhibitory neurons. We fasted mice the day before the experiment. Before starting, we removed every object from their homecage and connected mice to the patch cord. We waited 5 minutes to let the mice habituate before starting the recording. We recorded activity levels for 5 minutes as baseline and then placed a bowl with 1 mL ensure in one corner of the cage. The mice approached the bowl at their time, and we recorded the moment of the first lick, afterwards, we kept recording for 5 more minutes.

### Experimental timelines for InsCtx manipulation experiments (see below for further details)

#### Chemogenetic cohorts (CaMKII and VGAT-Cre)

##### Weeks 1-2

Habituation to scruffing, tail milking, experimental room and experimenter.

##### Week 3

Days 1-4: Ensure in homecage overnight. Same habituation as weeks 1-2
Day 4: Overnight fast
Day 5: Experimental day. Saline or C21 injection (approximately half for each condition)

##### Week 4

Days 1-4: Ensure in homecage overnight. Same habituation as weeks 1-2 Day 4: Overnight fast
Day 5: Experimental day. C21 or saline injection (alternate condition of the previous week)

#### Optogenetic cohorts

We created two cohorts: The first cohort (n_Chrimson_=8; n_tdtomato_=9) was used to test the effects of optogenetic inhibition on anticipatory insulin release, as well as post-meal insulin levels and glucose clearance (Fig. 3D-G). The second cohort (n_Chrimson_=11; n_tdtomato_=17) was intended for measurements that cannot be repeated within subject as they require large volumes of blood and harvesting tissues for biochemical analysis (Fig. 3H-K)

##### Weeks 1-2

Habituation to scruffing, tail milking, open cage, experimental room and experimenter

##### Week 3

Days 1-4: Ensure in homecage overnight. Same habituation as weeks 1-2 Day 4: Overnight fast
Day 5: Experimental day. No light turned on during anticipatory phase

##### Week 4

Days 1-4: Ensure in homecage overnight. Same habituation as weeks 1-2 Day 4: Overnight fast
Day 5: Experimental day. Light turned on 15 seconds before presenting ensure and until after blood sample was collected.

##### Week 5 (to control for order of experimental conditions)

Days 1-4: Ensure in homecage overnight. Same habituation as weeks 1-2 Day 4: Overnight fast
Day 5: Experimental day. No light turned on during anticipatory phase

#### Habituation for blood sampling

Each procedure took place in a quiet room with minimal to no presence of anyone except the experimenter. *Habitation for chemogenetic inhibition:* Mice started habituation 2 weeks after surgery unless signs of suffering or stress were detected. The first day of habituation involved 3 sessions separated by 2 hours of scruffing the mice 3 times, holding them for 5-6 seconds and releasing them into the home cage. The second day was the same except we mimicked the action of tail blood collection (“tail milking”) each time we scruffed them. Day 3, 4, 5 and 6 were the same as day 2 except with each scruffing, we added a sham intraperitoneal injection (without puncturing the skin) before mimicking the tail milking.

#### Different days of experience with Ensure

We habituated all mice to the empty bowl that we used for Ensure in their home cage for a few days before the experiments. We used an across-subject design due to physiological (and IACUC) restrictions of amount of blood withdrawal per week. Naïve mice were fasted overnight and received Ensure for the first time during the blood sampling protocol. For 1 and 2 days of exposure mice were fasted overnight and kept on mild food restriction (90-95% free feeding bodyweight) until the corresponding day of blood sampling, receiving 1 mL of Ensure per day (for the experience with Ensure). For the 4 days of exposure, mice received 1 mL Ensure in their homecage every day and fasted overnight before the blood sampling day.

#### Artificial sweetener experiment

We prepared a milkshake-like solution with ∼500 mM sucrose (480-540 mM, Sigma), ∼500 mK glucose (480-540 mM, Sigma), 2-2.25% Intralipid (Sigma) and 10-20% standard baking vanilla extract (Maimon’s), all diluted in water. Similar to our protocol with Ensure, we gave mice 1 mL per day of this milkshake-like mix in their homecage for the four days previous to the blood sampling. On the day of blood sampling, we gave mice a sugarless version of this solution with matched sweetness^83,84^, by replacing sucrose and glucose with acesulfame potassium (20-22.5 mM, Sigma) sucralose (0.8-0.9 mM, Sigma).

*Role of InsCtx in forming associations (learning) vs. expression of anticipatory insulin release:*

We food-restricted mice throughout the week of testing for the ‘learning’ portion of the experiment, keeping their weights between 85-90% of free feeding bodyweight. We gave mice 1 mL of Ensure daily and extra chow to maintain the target body weight. Ten minutes before giving Ensure, mice received an intraperitoneal C21 injection (3 mg/kg) and 2 hours later, a second C21 injection (1 mg/kg) to maintain InsCtx inhibition throughout the potential post-meal effects of Ensure. We then tested their anticipatory insulin response to test if InsCtx activity is necessary for associating the sensory properties of Ensure (e.g., sight, smell, taste) with its post-absorptive consequences (nutrient absorption). The week after the test, we gave mice 1 mL Ensure every day without food restriction and no C21 injection before testing the ‘expression’ portion of the experiment. That is, whether InsCtx activity is needed to produce an anticipatory insulin response when expecting Ensure.

*Habituation for optogenetic inhibition:* Mice started habituation 2 weeks after surgery unless signs of post-surgical suffering or stress were detected. The first day of the first week of habituation involved 3 sessions separated by 2 hours of scruffing the mice 3 times, holding them for 5-6 seconds and releasing them into the home cage. The second day was the same except we mimicked the action tail blood collection (“tail milking”) each time we scruffed them. Day 3 to day 5, involved the same scruffing habituation but incorporating time in an open cage for 15 minutes, 30 minutes and 45 minutes on day 3,4 and 5, respectively, after the first scruffing of each session and then two more scruffings afterwards. The “open cage” was the mice’s own home cage but with added high walls, with no access to water. We repeated this twice each day. At the end of day 2,4 and 5, a petri dish with 0.3mL Ensure (Vanilla, Abbot) was given to the mice in their homecage inside the animal facility to familiarize them with Ensure. On the next week, day 1 was the same as day 5 but optogenetic patch-cords were connected during the home cage habituation. This was repeated the rest of the week until the day of blood sampling.

*Pilot tissue harvesting:* Protocol was similar to the chemogenetic protocol but without sham intraperitoneal injections, and performed for 2 weeks.

*Tissue harvesting:* Protocol was similar to the optogenetic inhibition protocol but added ambient pulsing red light and 5mW blue light connected to the cannulas, in random time intervals in randomized order of 30, 45 and 60 seconds.

##### Blood sampling protocols

Mice were fasted overnight (14-18 hrs) the day before the blood sampling. On the day of the experiment, after taking the mice to the experimental room, mice were allowed to acclimate at least half an hour before starting any protocol.

Due to IACUC restrictions on the maximal volume of blood we can draw, C21/saline or light/no-light days occurred on different consecutive weeks.

*Chemogenetic cohorts:* We counterbalanced whether mice first had the control condition (saline) or the experimental condition (C21).

*Optogenetic experiments:* we analyzed 3 consecutive weeks (weeks 3-5 in the timelines above): no-light1◊light-ON◊no-light2 to show that the order of experimental conditions does not affect the anticipatory insulin responses (Supp. Fig. 3E).

##### Blood sampling for chemogenetic experiments

At the beginning of the experiment, we cut the tip of the mice’s tail and gave them an intraperitoneal injection with either saline (week 1) or Compound 21 (3 mg/kg, DREADD Agonist 21 dihydrochloride, Sigma Aldrich, week 2) in similar volumes and returned the mice to their home cage. After 10 minutes, we took the first blood sample for baseline levels of insulin and glucose by scruffing the mice and milking the tail into an anticoagulant collection tube (MiniCollect®, Greiner 1mL K3EDTA). After another 10 minutes, we placed a small Pyrex dish with 1 mL Ensure in the homecage. After the mice approached the petri dish, we ensured they licked the ensure at least once, immediately took the petri dish and waited 1 minute. The dish was placed near the cage for the mice to continue smelling and seeing the Ensure. After 1 minute, we took the second blood sample from the tail tip for anticipatory insulin and glucose levels. We placed the dish again in the cage and allowed mice to consume it freely for 10 minutes before taking the third blood sample from the tail tip for post absorptive insulin and glucose levels. Finally, we took the mice back to the animal facility with *ad libitum* access to food.

##### Blood sampling for optogenetic experiments

At the beginning of the experiment, we cut around 3mm from the tip of the tail of each mouse, waited 10 minutes, and then took the first blood sample for baseline levels of insulin and glucose by scruffing the mice and milking the tail into an anticoagulant collection tube. Afterwards, we connected the optogenetic patchcords to the optogenetic cannulas and left the mice for an hour in an open cage (their own home cage, see details above). In the optogenetic inhibition day, we then turned on the LED light (∼625 nm, 10 mW to each hemisphere; 10Hz, at 20% duty cycle; Orange-red LED, Prizmatix), waited 15 seconds and placed a bowl with 1 mL Ensure in their cages. In the control days (no optogenetic inhibition, No light 1 (week3) and No light 2 (week 5)), we directly placed the bowl with Ensure in the home cage. We allowed the mice to freely approach the bowl and waited until they took their first lick before immediately taking away the bowl and waiting 3 minutes. The dish was placed near the cage to allow the mice to continue smelling and seeing the Ensure. After 3 minutes, we took the second blood sample from the tail tip for anticipatory insulin and glucose levels. Next, we placed the Ensure bowl again in the cage and allowed mice to consume it freely for 10 minutes before taking the third blood sample from the tail tip for post absorptive insulin and glucose levels. Five minutes, 15 minutes and 25 minutes later, we measured glucose levels from the tail tip with a FreeStyle Optium Neo glucometer (Abbott) to complete the glucose curve. Finally, we took the mice back to the animal facility with *ad libitum* access to food.

In our earlier chemogenetic experiments we sampled cephalic insulin 1 min after Ensure presentation. However, consistent with previously published work showing stronger cephalic phase insulin levels 3 min vs. 1 min from consumption onset^45^, in subsequent experiments we noticed that 3 min post Ensure presentation tended to yield more consistent cephalic phase insulin levels than 1 min post Ensure presentation. We therefore chose to use 3 min for our optogenetic experiments. We note that the aforementioned study by Glendinning et al.^45^ did allow mice to consume in this 3 min time interval, while we did not. Thus, it is also highly unlikely that mice in our study consumed enough Ensure in one lick to drive a sufficient rise in blood glucose levels that would drive post-absorptive glucose-dependent insulin release.

##### Blood sampling from head-fixed mice

We trained mice to consume Ensure from a lickspout while head-fixed, similar to conditions during two-photon imaging during Ensure refeeding (see below). We food-restricted mice to consume large quantities of Ensure (multiple consecutive drops, each drop ∼5 µL, 1 drop every 2 sec). We initially sampled blood from the tail tip as described above for baseline levels. Ten minutes later, we allowed mice to consume 2 drops of Ensure (total ∼10 µL), waited 2 min, and then sampled blood again for cephalic phase levels.

##### Blood sampling for protocol food type selection

Same as above chemogenetic protocol. Food was either 1 or 10 14 mg pellets (Bioserv), 4g of Chow, 1mL Ensure, or 4g of Chocolate (Alpine milk, Milka). We sampled blood immediately after, 1 minute after or three minutes after food presentation.

##### Blood sampling for insulin kinetics

Same as chemogenetic protocol. However, mice had a waiting period of 1, 3 or 7 minutes after presentation and first lick of ensure, before taking the blood sample for anticipatory levels. Each mouse went through each waiting period on different days.

##### Blood sampling with artificial sweetener mix

Same as chemogenetic protocol but mice received a new sugarless milkshake-like solution (see details above). After the presentation of the artificial sweetener mix and first lick, we waited 1 minute before blood collection.

In all cases, the blood collection tubes were kept on ice until centrifugation. The glucose levels of the first three samples were measured by taking approximately 1.2 µL from the blood collection tube and placing it on the glucometer strip.

Blood samples were centrifuged (Eppendorf, Centrifuge 5425 R) at 2000 rpm for 20 minutes at 4°C. Then the supernatant (plasma) was transferred to PCR tubes before freezing at −80°C until further analysis. For the heart puncture blood samples, they were left at room temperature for 30 minutes to two hours before centrifuging for 10 at 2000rpm, at 4C. We collected the supernatant (serum) and froze samples at −80C until analysis.

#### Tissue collection (Pilot and optogenetic inhibition protocol)

On tissue harvest days, we followed the same optogenetic protocol except that after 10 minutes of free Ensure consumption, we waited 5, 10 or 15 minutes (for the pilot) or 5 minutes (optogenetic inhibition protocol) before using isoflurane for terminal anesthesia. We then collected blood by heart puncture and extracted the liver. We snap-froze all samples in liquid nitrogen and kept them at −80° C until processing.

#### Assessment of the relationship between cephalic insulin levels and time-of-day

To evaluate the relationship between cephalic phase insulin release and potential circadian regulation, we calculated the correlation between cephalic phase insulin release and the time-of-day of blood collection.

Both baseline insulin levels and normalized cephalic phase levels were not significantly correlated with time-of-day. We also binned the time of day into four segments to further show that there is no relationship in our protocol between time-of-day and circulating insulin levels or anticipatory insulin released.

#### Inclusion of non-responders

There is a well-known phenomenon in the literature of “physiological non-responders” that lack, or have very low, anticipatory insulin release in mice, rats, and humans^39,51,85^. This has been shown to be a stable feature of individual mice across multiple measurements^50^. Therefore, we excluded mice from analyses if they exhibited a negative change in insulin in *<u>control conditions</u>* (saline injection day for chemogenetics and no-light day for optogenetics). We then removed them from <u>both</u> analyses: the control day, and their corresponding experimental day (C21 injection or light-on). Therefore, all the mice we did include in analyses actually may exhibit a very small or even approximately zero change in insulin and still be included in the analysis. Importantly, we verified that including non-responders in analyses does not affect any of the results in both chemgenetic and optogenetic experiments (Supp. Figs. 1G, 3E, 4A,B).

Some of these non-responder mice from the optogenetic across-subject cohorts (n_ChRimson_=5; n_tdTomato_=3) were then used for the analysis of the effect of the optogenetic protocol on blood glucose in the absence of elevated anticipatory insulin (Supp. Fig. 3J,K), as well as a separate cohort not included in this manuscript due to technical issues with AAV expression (n=7).

One extreme outlier, with insulin values consistently >2.5 STD above population mean of baseline insulin levels on multiple days, irrespective of experimental conditions, was also removed from analysis.

##### Biochemistry measurements

Insulin levels were determined from plasma with an ultrasensitive mouse insulin ELISA kit (#90080, Crystal Chem). We followed the kit instructions except that the first 2 hours incubation was on a shaker at 4°C and the following incubation on a shaker at room temperature. We measured free fatty acids from serum samples with the LabAssay (TM) NEFA-HR (633-52001, FUJIFILM Wako)

##### Brain tissue preparation and immunohistochemistry

We terminally anesthetized mice with Pentobarbital (Pental 20%, Vetmarket) and transcardially perfused with phosphate-buffered saline (PBS) followed by exclude neutral buffered Formalin (Sigma Aldrich). We then extracted the brains and put them in a cryoprotection solution of 30% sucrose in PBS. The next day, we sectioned them coronally on a freezing sliding microtome (Leica Biosystems) with 50 μm thickness, collected in 3 equal series.

We washed the brain sections in PBS, added blocking solution (3% fetal bovine serum/0.25% Triton X-100 in PBS) for 1 hour at room temperature in a shaker and then incubated the samples overnight at room temperature in blocking solution containing primary antiserum (1:1000, Living Colors dsRed Polyclonal Antibody, Takara or 1:500 anti-GAD1/2, Abcam, AB183999). The next morning, we washed the sections 3 times in PBS for ten minutes on a shaker and then incubated in Alexa fluorophore-conjugated secondary antibody (Molecular Probes, 1:1000, ThermoFischer) for 2 h at room temperature. After 3 more 10 minutes washes in PBS, sections were mounted onto microscope slides and fluorescent images were captured with an Olympus BX61VS slide scanner microscope at 10X. All antibodies used were previously verified.

#### Histological verification

Mice with incorrect virus expression or fiber targeting were excluded from all experimental analysis. Reasons for exclusion included: Absence of viral expression or discernible fiber tract in one of the two hemispheres, viral expression or fiber tract outside target area, or undetermined fiber tract. Histology was analyzed by a blinded investigator to prevent bias. Nevertheless, in some mice we observed some viral expression that also spread beyond InsCtx into the adjacent secondary somatosensory cortex (dorsally) and piriform cortex (ventrally). We therefore analyzed the effect of this viral spread outside the InsCtx to confirm that the inhibitory effects observed both from chemogenetic and optogenetic inhibition are due to effects on InsCtx activity and not from an effect on other cortices. To this end, we sub-selected mice from the chemogenetic cohort that had viral spread only in InsCtx and found similar extent of inhibition compared to the whole group. Saline: 0.67 ±0.13 vs. C21: −0.08±0.1 (mean ± SEM), p=0.002, n=5. This suggests that the effect seen from the whole cohort is due to the inhibition on the InsCtx and not from non-specific effects over other cortices. Moreover, in optogenetic inhibition experiments, we verified that all mice had fiber tips scars in the InsCtx, thereby precluding effects on the dorsally adjacent secondary somatosensory cortex. We also conducted a similar analysis to control for spread ventrally to piriform cortex and observed similar results: no-light: 1.39 ±0.169 vs. light: 0.16±0.2 (mean ± SEM), p=0.03, n=6.

##### GAD1/2 imaging and quantification

We imaged InsCtx slices using a Thorlabs Bergamo two-photon microscope via a 10X objective (TL10X-2P) with a resolution of 2048x2048 pixels (pixel size 0.3 µm), separately using 920 nm excitation (FFUltra 920, Toptica) for the green channel (anti-GAD), and 1050 nm excitation (FFUltra 1050, Toptica) for the red channel (mCherry or tdTomato). We quantified red/green overlap semi-manually using CellPose^86^.

##### Western blots

Pulverized liver samples (∼50mg) were homogenized in 500ul RIPA buffer containing proteases inhibitor cocktail. Samples were cleared using centrifugation at 18,000g at 4C (Centrifuge 5427 R, Eppendorf) and 10 µg of total protein were used for western blot analysis. and separated the supernatant. We next incubated the membrane with primary antibodies 1:1000 in 5%BSA. Anti-phosphorylated Akt in S473 (Rabbit, Cell Signaling technology #4060), anti-phosphorylated GSK3b in S9 (Rabbit, Cell Signaling technology #5558) and anti-Vinculin (Rabbit, Cell Signaling technology #13901). Incubation was overnight on a shaker at 4C. The next day we washed membrane 3 times for 10 minutes in shaker at room temperature with TBST (10%TBS, 0.1% Tween). We next incubated with secondary antibody anti rabbit (1:10000m Jackson ImmunoResearch) in 5% skim milk (Difco Skim Milk, BD) in TBST for 2 hours in shaker at room temperature. We washed the membranes again with TBST 3 times for 15 minutes in shaker at room temperature. We prepared the solution for the electrochemiluminescence reaction with equal parts of reagents A and B of the ECL Prime Peroxide Solution kit (Cytiva). We imaged the membranes in the ChemiDoc Imaging system (BIO-RAD). Afterwards, membranes were stripped of the antibodies with an Antibody stripping buffer (Gene Bio-Application) for 10 minutes. We washed them 5 times for 5 minutes at room temperature on a shaker. We then repeated the process of incubation with primary anti-total protein antibodies: anti-total Akt (Mouse, Cell Signaling technology #2920) or GSK3b (Rabbit, Cell Signaling technology #12456), repeated the incubations, the secondary antibodies anti-rabbit or mouse (1:1000, Jackson ImmunoResearch) in 5% skim milk, and the imaging process.

Analysis of the membranes was done with ImageLab Software (BIO-RAD) to measure band intensity and then calculate the ratio of the phosphorylated band volume over the total protein band volume.

##### Fiber photometry data analysis

Traces sampled at 45 or 50 Hz and smoothed using a 2.2s sliding running average. For each channel separately, we first denoised the data with a median filter, and then corrected for bleaching by subtracting a double exponential fit of the data. We then calculated the difference between channel 1 (465 nm) and 2 (405 nm, approximate isosbestic point) and z-scored the data. z-score = (F – mean F0)/ std F0. F0 is the average of the first 5 minutes before the saline injection or before the LED was turned on. For fiber photometry analysis of excitatory projection neurons or inhibitory interneurons while mice approach Ensure, we aligned all traces to the first lick before averaging them and re-zeroed the z-scored data based on the mean of the 20 sec before the first lick.

##### 2-photon imaging during refeeding

Two-photon imaging of GCaMP6f was performed using a resonant-scanning two-photon microscope with tiltable scanhead (Neurolabware; 31 frames/second; 1154x512 pixels). All imaging was performed with a 20x 0.45 NA air objective (Olympus) with a 540 x 360 μm2 field of view. All imaged fields of view (FOV) were at a depth of 90-200 µm below the pial surface, using a Mai Tai DeepSee laser (Newport Corp.) with laser power at 920-960 nm of 35-80 mW at the front aperture of the objective (power at the sample was likely substantially less due to partial transmission via the microprism). Imaging depth was adjusted in between runs (every 30 min) to account for slow drift in the z plane (< 7 μm). Recording locations were approximately +0.5 mm to −1.0 mm to the middle cerebral artery (see refs. ^12,15^ for further details).

We initially imaged fasted mice while they performed a short 30 min session of the visual cue discrimination task^12,15^. After this 30-min session, in which they consumed ∼0.3 mL Ensure, we started the refeeding session (Fig. 2, and Supp. Fig. 2). Importantly, we and others have previously shown that mice are still highly motivated to consume Ensure at this phase^12,15,87^. We initially imaged baseline activity for ∼30 sec and delivered 1 drop of Ensure (∼5 µL) thereby providing initial olfactory cues, which then drive consumption. We thus followed InsCtx activity before Ensure consumption, and as mice began consuming Ensure ad libitum until voluntary cessation of consumption. Ensure consumption lasted 60-90 minutes. We triggered delivery of Ensure with every lick, but with a minimum interval of 2 sec between Ensure deliveries. During this period of time, mice consumed ∼3-5 mL of Ensure and then voluntarily stopped licking for Ensure. We then imaged mice for an additional 30 min post refeeding while they were presented with visual cues but did not overtly respond to them^12,15,87^. Ongoing activity in inter-trial intervals in this last session served as the “refed state”. Importantly, we have previously shown that in such task contexts, ongoing activity patterns in InsCtx during inter-trial intervals faithfully reflect the current physiological state^12^.

##### 2-photon data analysis

Each acquired image was spatially down sampled by 2X. To correct for motion along the imaged plane (x-y motion), each frame was registered to an average field-of-view using efficient subpixel registration methods^88^. Within each imaging session, each run (2-8 runs/session) was registered to the first run of the day. Image stacks were de-noised using principal components analyses (PCA) of every pixel across time, and by user identification and removal of noise principal components (low eigenvalues). Cell masks and calcium activity time courses (‘F(t)’) were extracted using custom implementation of common methods^89^. To avoid use of cell masks with overlapping pixels, we only included the top 75% of pixel weights for a given mask, but users screened each prospective ROI and could edit the size of the mask, selectively removing the lowest probability pixels. We then excluded any remaining pixels identified in multiple masks. We manually verified that all cell masks had typical cell body morphology and size.

Fluorescence time courses were extracted by averaging the pixels within each region-of-interest (‘ROI’) mask. Fluorescence time courses for neuropil within a 25 µm annulus surrounding each ROI (but excluding adjacent ROIs and a protected ring surrounding each ROI) were also extracted (Fneuropil(t): median value from the neuropil ring on each frame). Fluorescence timecourses were calculated as Fneuropil_corrected(t) = FROI(t) - Fneuropil(t). The change in fluorescence was calculated by subtracting a running estimate of baseline fluorescence (F0(t)) from Fneuropil_corrected(t), then dividing by F0(t): ΔF/F(t) = (Fneuropil_corrected(t) - F0(t))/ F0(t). F0(t) was estimated as the 10th percentile of a 32 sec sliding window ^12,15,87^.

We z-scored neuronal activity based on the pre-consumption baseline period. For evaluation of the initial burst of activity before and at meal onset, we categorized neurons as having increased or decreased activity if their activity z-score during the 4 sec before consumption onset or during the first min post meal onset was above 1.5 or below −1.5. We calculated response magnitude at the average z-scored activity during these time windows.

For presentation purposes, all licking, mean neural activity, and projected neural activity were smoothed with a 10 sec leading edge running average. To project InsCtx population activity patterns onto the axis of fasted vs. refed activity^12^, we defined this axis for each mouse by 𝑥̅*_fasted_* − 𝑥̅_*refed*_, where 𝑥̅ is the population vector of mean activity in a given state. For the fasted state this was the mean activity in the baseline period until 4 sec before refeeding, while in the fed state, this was the mean activity in the refed session. We projected activity onto this axis by calculating the dot product of this vector with the time-varying pattern of InsCtx population activity, x(t). We then scaled values along this axis per mouse, ascribing a value of 1 when x(t) =𝑥̅*_fasted_* and value of 0 when x(t) = 𝑥̅_*refed*_ , and intermediate values for patterns that fall between 𝑥̅*_fasted_* and 𝑥̅_*refed*_.

We sought to verify that the changes we observed during the onset of refeeding truly reflected a specific *pattern* of activity and were not simply due to overall increase in neural activity across neurons (Supp. Fig. 2D, although note that many neurons also decreased activity, Supp. Fig. 2A). To this end, for each mouse we artificially multiplied baseline neural activity by a factor of 2 and then brought it back to baseline, thus mimicking the mean activity profile. In essence, this increased population vector lengths but without changing vector direction (the pattern).

Two-photon imaging data used for this manuscript were not previously analyzed or published. These data were acquired in the context of larger longer-term experiments examining InsCtx activity in different feeding contexts, some of these data (not in the current manuscript) were previously published^12,12,53^.

##### Miniscope imaging during refeeding

Animals were habituated to liquid food (Ensure Plus, CVS) by putting 0.3 mL in a 60 mm-diameter petri dish lid in home cage at the beginning of the dark cycle. After 4 days of habituation, mice were fasted overnight. During the imaging experiment, mice first habituated by placing them in clean home cages, connecting the miniscope was connected waiting 5 minutes. We then recorded 5 minutes of baseline ongoing activity. We then added 1 mL of Ensure Plus to the petri dish and continued recording for 10 min after mice began consumption. Next, we added 3 mL Ensure Plus to the petri dish, and waited for 1 hour, after which Ensure Plus was then removed and measured. Finally, we imaged 10 minutes of ongoing activity after Ensure Plus removal (refed session).

#### Miniscope imaging and behavioral data analysis

Calcium imaging data was collected by an Inscopix DAQ box with 20 Hz acquisition speed and preprocessed by Inscopix IDPS software. Raw imaging data was reduced to 10 Hz from 20 Hz, then ΔF/F values were calculated. The baseline of fluorescence was the mean of entire video. Individual cells were first obtained using the PCA/ICA approach, then were manually accepted or rejected.

We z-scored neuronal activity based on the 5 min pre-consumption baseline period. For evaluation of the initial burst of activity before and at meal onset, we categorized neurons as having increased or decreased activity if their activity z-score during the 10 sec before consumption onset or during the first 30 sec post meal onset was above 1.5 or below −1.5. We calculated response magnitude at the average z-scored activity during these time windows.

For presentation purposes mean neural activity, and projected neural activity were smoothed with a 30 sec leading edge running average. To project InsCtx population activity patterns onto the axis of fasted vs. refed activity ^12^, we defined this axis for each mouse by 𝑥̅*_fasted_* − 𝑥̅_*refed*_, where 𝑥̅ is the population vector of mean activity in a given state. For the fasted state this was the mean activity in the baseline period until 5 min before refeeding, while in the fed state, this was the mean activity in the refed session. We projected activity onto this axis by calculating the dot product of this vector with the time-varying pattern of InsCtx population activity, x(t). We then scaled values along this axis per mouse, ascribing a value of 1 when x(t) =𝑥̅*_fasted_* and value of 0 when x(t) = 𝑥̅_*refed*_, and intermediate values for patterns that fall between 𝑥̅*_fasted_* and 𝑥̅_*refed*_.

### Phosphoproteomics

#### Sample preparation

Lysates were sonicated six cycles of 30 s (Bioruptor Pico, Diagenode, USA). Protein concentration was measured using the BCA assay (Thermo Scientific, USA) and a total of 120 μg protein was reduced with 5 mM dithiothreitol (Sigma) and alkylated with 10 mM iodoacetamide (Sigma) in the dark. Each sample was loaded onto S-Trap minicolumns (Protifi, USA) according to the manufacturer’s instructions. In brief, after loading, samples were washed with 90:10% methanol/50 mM ammonium bicarbonate. Samples were then digested with trypsin (1:50 trypsin/protein) for 1.5 h at 47 °C. The digested peptides were eluted using 50 mM ammonium bicarbonate; trypsin was added to this fraction and incubated overnight at 37 °C. Two more elutions were made using 0.2% formic acid and 0.2% formic acid in 50% acetonitrile. The three elutions were pooled together and vacuum-centrifuged to dry. Samples were kept at −20 °C until analysis^90^.

#### Immobilized Metal Affinity Chromatography

115 μg of each sample was subjected to phosphopeptide enrichment. It was performed on a Bravo robot (Agilent Technologies) using AssayMAP Fe(III)-NTA, 5 μL cartridges (Agilent Technologies), according to the manufacturer’s instructions. In brief, cartridges were primed and equilibrated with 50 μL of buffer A (99.9% ACN/0.1% TFA) and 100 μL if buffer C (80% ACN/19.9% H2O/0.1% TFA), followed by sample loading in 100 μL of buffer C at 5 μL/min. Phosphopeptides were eluted with 120 μL of buffer B (99% H2O/1% NH3) at 5 μL/min. Three μL of formic acid was added to each sample for acidification. Prior to LC–MS analysis, all samples were dried down to the volume of 15 μL^90^.

#### Liquid chromatography

ULC/MS grade solvents were used for all chromatographic steps. Each sample was loaded using split-less nano-Ultra Performance Liquid Chromatography (Acquity M Class; Waters, Milford, MA, USA). The mobile phase was: A H2O + 0.1% formic acid and B acetonitrile + 0.1% formic acid. Desalting of the samples was performed online using a reversed-phase Symmetry C18 trapping column (180 µm internal diameter, 20 mm length, 5 µm particle size; Waters). The peptides were then separated using a T3 HSS nano-column (75 µm internal diameter, 250 mm length, 1.8 µm particle size; Waters) at 0.35 µL/min. Peptides were eluted from the column into the mass spectrometer using the following gradient: 4% to 20%B in 115 min, then to 90%B in 7 min, maintained at 90% for 7 min and then back to initial conditions.

#### Mass Spectrometry

The nanoUPLC was coupled online through a nanoESI emitter (10 μm tip; Fossil, Spain) to a quadrupole orbitrap mass spectrometer (Exploris 480, Thermo Scientific) using a FlexIon nanospray apparatus (Thermo). Data was acquired in data dependent acquisition (DDA) mode, using a ‘top-speed’ method, cycle time of 2sec. MS1 resolution was set to 120,000 (at 200 m/z), mass range of 380-1500 m/z, AGC of 200% and maximum injection time was set to 50 msec. MS2 resolution was set to 15,000, quadrupole isolation 1.4m/z, AGC of 150%, dynamic exclusion of 40 sec and maximum injection time of 120 msec.

#### Data processing

The data analysis was performed using MetaMorpheus version 1.0.2^91^, available at https://github.com/smith-chem-wisc/MetaMorpheus.

The following search settings were used: protease = trypsin; search for truncated proteins and proteolysis products = False; maximum missed cleavages = 2; minimum peptide length = 7; maximum peptide length = unspecified; initiator methionine behavior = Variable; fixed modifications = Carbamidomethyl on C, Carbamidomethyl on U; variable modifications = Oxidation on M; max mods per peptide = 2; max modification isoforms = 1024; precursor mass tolerance = ¬±5.0 PPM; product mass tolerance = ¬±20.0 PPM; report PSM ambiguity = True. The combined search database contained 17417 non-decoy protein entries including 476 contaminant sequences. We applied the G-PTMD module in MetaMorpheus to identified phosphorylation of S, T or Y in combination of other modifications (default list) ^92^. The LFQ intensities (Label-Free Quantification) were extracted and used for further calculations using Perseus v1.6.2.3. Decoy hits were filtered out. The data was further filtered to include only peptides with at least 3 valid values in at least one of the groups. Missing values were imputed by selection from a low random distribution (width=0.3, downshift=1.8). Analysis of Variance (ANOVA), after logarithmic transformation, was used to identify significant differences across the biological replica.

#### Permutation test for assessment of false positive rates

For each phosphosite, we shuffled the log2(fold change) of each subject and reassigned it to two new groups with either n=11 or n=17 to mimic the real sample sizes in our experimental and control groups. Then, we calculated the mean of each group and compared them with a Student’s t-test. We then removed phosphosites that had a log2 (fold change) between −0.5 and 0.5 and calculated for the remaining phosphosites, the adjusted p-value with the Benjamini-Hochberg correction. We repeated this for 1000 permutations. After that, we chose two target FDRs: 5% (similar to an α=0.05) and 15% (which is a standard in the phosphoproteomics field). For the histogram graph (Supp. Fig. 4J), we calculated the normalized frequency in percentage as the counts per bin over the total number of counts multiplied by 100. We divided the data into 50 bins.

#### Phosphoproteomics analysis

The volcano plot was generated using the uncorrected p-values to plot our complete data (including phosphosites with a log2(fold change) between −0.5 and 0.5). Using the Gene Ontology database (gene2go.gz, https://ftp.ncbi.nlm.nih.gov/gene/DATA/) to access all the GO terms available and its related genes (Table S1, spreadsheet: ‘GO_terms’), we generated ‘global terms’ related to different aspects of metabolism: Insulin signaling, lipid metabolism, carbohydrate metabolism and, separately, gluconeogenesis. We separated these last two in an effort to understand if we could attribute the difference in glucose clearance to either enhanced lipid metabolism or reduced gluconeogenesis. We searched the GO database for terms that would be related to each global term (See Table S2, spreadsheet: ‘metabolic_GO_terms’) and merged them by creating new lists of all genes belonging to each metabolism category and removing duplicated genes if necessary. The search of GO terms belonging to each category was done by finding specific words (See Table S2, spreadsheet ‘Criteria’). In some cases, the search included GO terms that were not specifically related to the global terms we built so we discarded them. For example, when looking for the word ‘metabolism’, a lot of GO terms related to DNA, RNA and aminoacids metabolism were highlighted but we considered them not relevant and discarded them.

We filtered our phosphosites by their adjusted p-value, keeping only p-values < 15% FDR (p<0.1315) and matched then to each global term, allowing matches with more than one term. This information was then plotted on top of the volcano plot, keeping the uncorrected p-value coordinates.

For both the significantly up-regulated vs down-regulated phosphosites plot (Supp. Fig. 4I) and the significant phosphosites in a global metabolism category (Fig. 3F), we used as ‘background’ the phosphosites with a log2(fold change) between −0.5 and 0.5 (Supp. Fig. 4J,K) for the Fischer’s exact test, doing the analysis with the Fischer exact Python package in scipy stats.

For the individual gene plots, we plotted each separately but used the adjusted p-values calculated as described above, all of them falling below 0.05 of the adjusted p-value which means less than 5% FDR according to our method.

### Statistics

Statistical tests were performed using standard Python and Matlab functions. We started by testing normality (Shapiro-Wilks test) and homogeneity of variance (Barlette’s test) of the sample distributions. If data were normal with homogenous variance we used parametric tests (Student’s independent or paired t-test). If we made multiple comparisons, we corrected with Benjamini-Hochberg procedure. If normality was not met, we used non-parametric tests (Mann-Whitney U or Wilcoxon signed rank test). For comparing more than two conditions we used ANOVA or repeated measures ANOVA (normal distribution) or Welch’s ANOVA (non-normal distribution). For correlation analysis, we used Pearson’s correlation coefficient (normal distribution) or Spearman’s rank test (non-normal distributions). For phosphoproteomics analysis, we followed the standards in the field and used an independent student t-test with adjusted p-values using the Benjamini-Hochberg FDR correction, and Fischer’s exact test for comparing proportions of phosphorylation events against background phosphorylation.

---

**Supp. Fig. 1:**
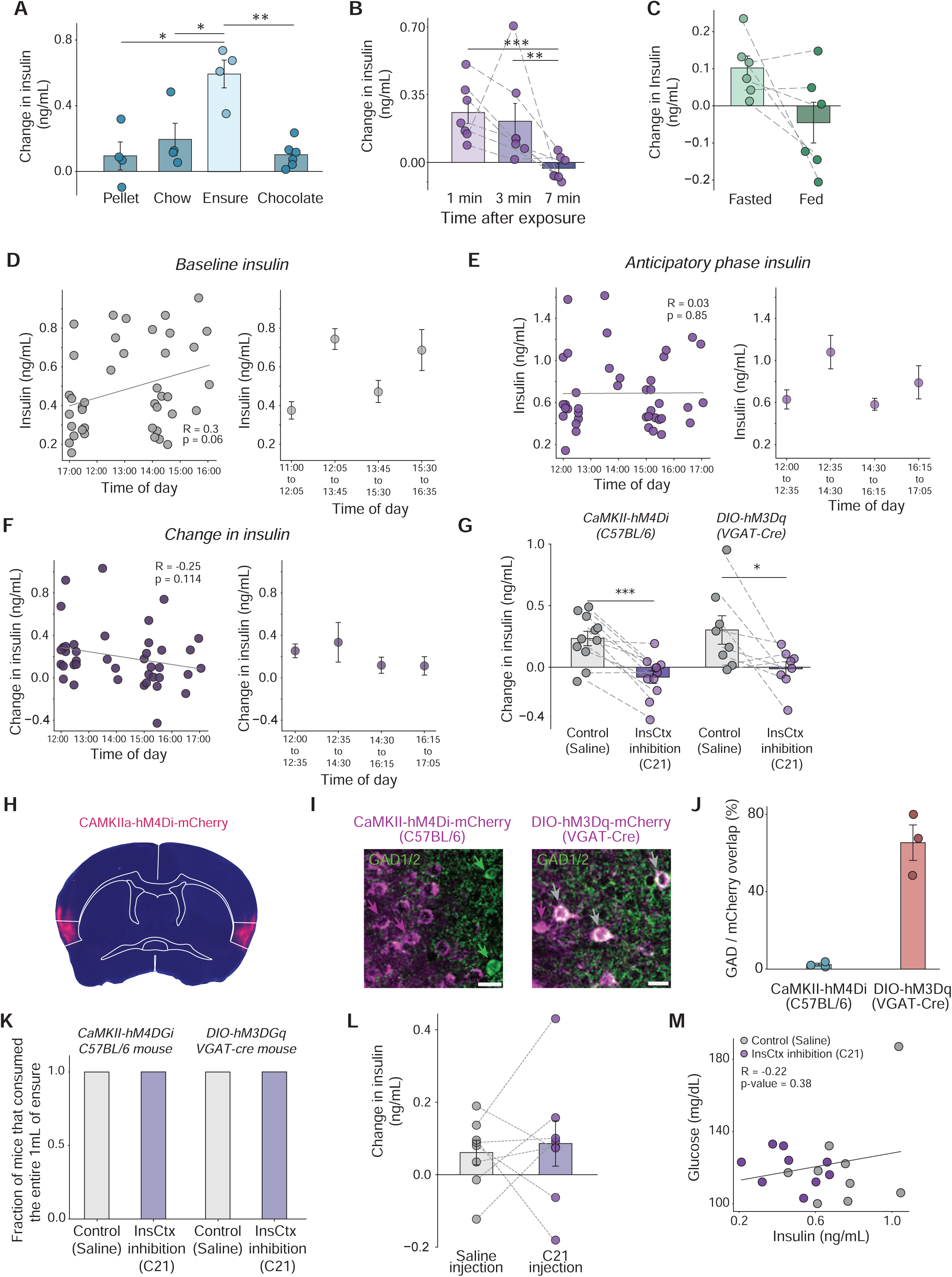
Additional experiments and analyses for the learned anticipatory insulin release assay and for chemogenetic inhibition experiments. **(A)** Comparison of anticipatory insulin release across 4 different types of food. Ensure was the food type that elicited the highest and most consistent anticipatory insulin release (n_Pellet, Chow, Ensure_=4, n_Chocolate_=6, p_Ensurevs.Pellet_=0.002, p_Ensurevs.Chow_=0.01, p_Ensurevs.Chocolate_=0.0009). **(B)** Anticipatory insulin kinetics. Normalized change in insulin levels 1, 3 or 7 minutes after presenting the bowl of Ensure during the protocol, mice had similar levels in the first 1-3 minutes (n=7, p=0.68), declining thereafter to baseline after no further exposure (p_1vs7_=0.002. P_3vs7_=0.04). **(C)** Overnight fasted mice (14 −18 hrs) had higher anticipatory insulin release compared to the same mice without fasting (n=6, p = 0.05). **(D-F)** Analysis of the effect of time-of-day on insulin levels. **(D)** No significant correlation between time-of-day and circulating baseline insulin levels in fasted mice, despite a positive trend (n=40, R= 0.3, p=0.06). **(E)** No significant correlation between time-of-day and anticipatory insulin release levels (n=40, R= 0.03, p=0.85). **(F)** No significant correlation between time-of-day and the change in insulin (n=40, R= 0-0.25, p=0.11). **(G)** Re-analysis of the two different approaches to chemogenically inhibit InsCtx, without removing non-responders, show similar significant effect in anticipatory insulin (see Methods; CaMKII-hM4Di: n=11, p=0.0001; VGAT-hM3Dq: n=8, p=0.02). **(H)** Example of coronal slice at bregma with bilateral hM4Di-mCherry expression. Scale bar: 1 mm. **(I)** Example images of brains from slices at bregma showing specificity of viral expression. *Left*: AAV-CaMKII-hM4Di in C57BL/6 mice show very little overlap with anti-GAD1/2 immunostaining. *Right:* AAV-DIO-hM3Dq in a VGAT-Cre mic with relatively high overlap between virus expression and immunostaining. Magenta arrow: viral expression, green arrow: somatic anti-GAD1/2 labeling, grey arrows: neurons with overlap. Scale bar: 20 µm. **(J)** Quantification of the percentage of overlap between neurons expressing the virus and labeled with the anti-GAD1/2 antibody for both cohorts (green and red out of all red). n_CaMKII_= 104±22, n_VGAT_= 114±5 neurons from 3 mice each. **(K)** All mice, from both cohorts, consumed the total amount of Ensure given (1 mL) in both experimental days (n_CaMKII_=9, n_VGAT_=7). **(L)** Control experiment for chemogenetic inhibition showing that C21 injection in mice expressing a non-chemogenetic virus in the InsCtx did not reduce anticipatory insulin release (n=8, 0.73). **(M)** No significant correlation between glucose levels and insulin levels during the anticipatory phase (n=9, R = 0.081, p = 0.74), indicating that the increase in anticipatory insulin release in the control day is not triggered by increases blood glucose). Data are presented as individual measurements (points) and mean (bars) ± SEM. Gray dashed lines represent the same mouse. P values were calculated by Student’s paired two sample t-test (for **C,G,L**), Pearson correlation (for m), Spearman’s rank test (for **D-F**), ANOVA with post-hoc Tukey test (for **A**), or repeated measures ANOVA with pairwise post-hoc test Benjamini-Hochberg (for **B**). * p<0.05, ** p<0.01.

**Supp. Fig. 2:**
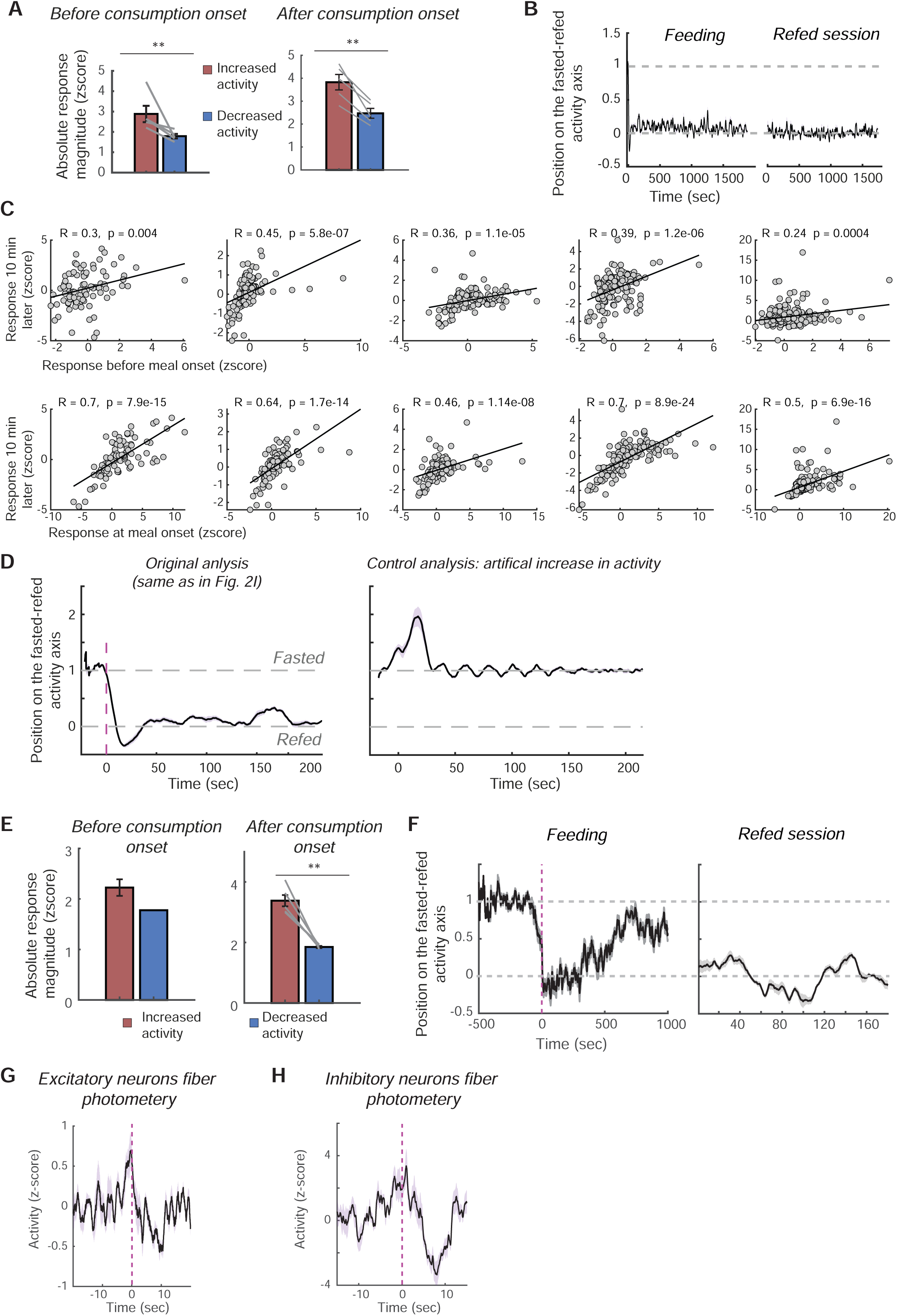
Additional analyses for InsCtx two-photon imaging and one-photon miniscope imaging experiments. **(A)** Absolute response magnitude of increased vs. decreased responses before (p = 0.03) and at (p = 0.002) meal onset. **(B)** Position on the fasted-refed axis of population activity shown for the entire 30 min feeding session and the entire 30 min refed (post-meal) session. Note that activity remains stable around values of 0 in the refed session. **(C)** Correlation across neurons per mouse between the activity before onset (top) or at meal onset (bottom) and the activity 10 min later. Note significant and high correlations but with substantially lower values 10 min later. Each plot column is from a different mouse. **(D)** Control analysis for the specificity of InsCtx meal onset activity patterns. We artificially increased activity by multiplying the baseline activity level of each neuron by a factor of 2. This resulted in an opposite effect, suggesting the burst of activity at meal onset is a specific pattern, distinct from the baseline (fasted) pattern, regardless of overall activity levels. **(E)** Same as ‘A’ but for miniscope data. **(F)** Same as ‘B’ but for miniscope data. **(G-H)** Bulk activity fiber phototmetry from excitatory neurons (**G**) and inhibitory (**H**). Vertical dashed line: licking onset. n= 4 mice for each group. Values are Mean ± SEM across mice. The p-values were calculeted by Student’s paired t-test. * p<0.05, ** p<0.01

**Supp. Fig. 3:**
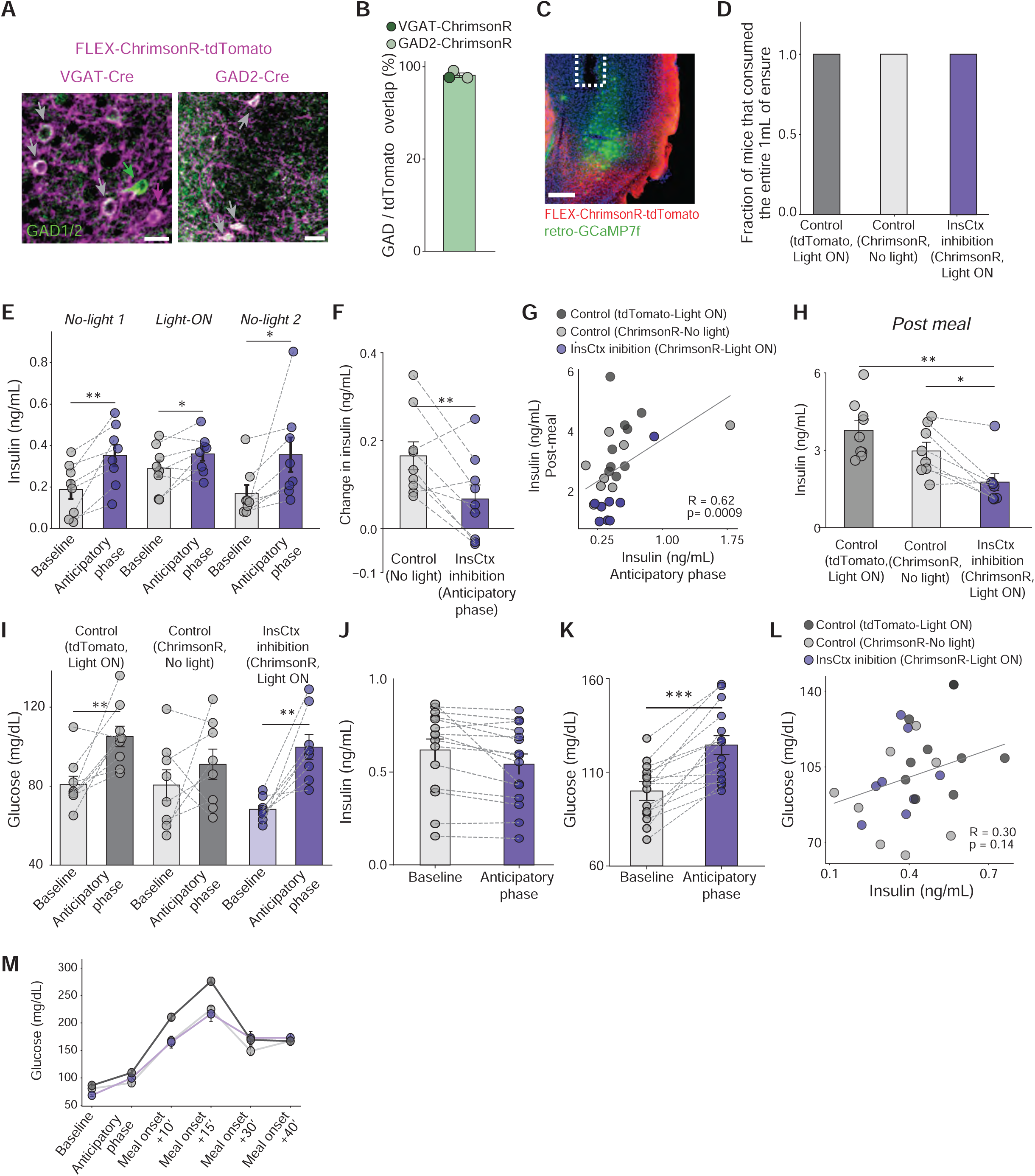
**Further control experiments and analyses for within-subject optogenetic inhibition experiments.** (**A**) Example images from brains slices at bregma showing specificity of viral expression. *Left*: AAV-FLEX-ChrimsonR in a VGAT-Cre mice shows high overlap with anti-GAD1/2 immunostaining. *Right:* AAV-FLEX-ChrimsonR in a GAD2-Cre mice with high percentage of overlap between virus expression and the antibody. Green arrow: anti-GAD1/2, grey arrows: neurons with overlap. Scale: 20 µm. (**B**) Quantification of the percentage of overlap between neurons expressing the AAV and labeled with the anti-GAD1/2 antibody for both Cre lines shows high specificity of viral expression in interneurons. *Dark green*: VGAT-Cre, *light green*: GAD2-Cre n= 83 ±30 (mean ±SEM) neurons from 3 mice. (**C**) Example section with co-expression of GCaMP7f (in excitatory projection neurons) and ChrimsonR (in inhibitory interneurons), and fiber placement. Scale bar: 200 μm. (**D**) All mice consumed the total amount of Ensure given (1 mL) in all experimental days (n_tdTomato_=9, n_ChrimsonR_=8). (**E**) Control for the order experimental condition (Light ON or OFF) demonstrates that the order of conditions does not underlie the effects of optogenetic inhibition as mice show normal anticipatory insulin the following week with no light during the protocol. First week: mice had no light during the protocol (No Light 1) and normal anticipatory insulin release (n=8, p=0.004). Second week: we turned on LED during the anticipatory phase (Light ON) and mice had reduced anticipatory insulin release (p=0.044, see Fig. 3). Third week: no light during the anticipatory phase (No Light 2) and mice recovered their anticipatory insulin release (p=0.006). See Fig. 3 for insulin change values per condition. (**F**) Re-analysis of the effect of optogenetic inhibition of InsCtx on anticipatory insulin, without removing non-responders, demonstrates a similar effect (n=9, p=0.008). (**G**) Re-analysis of the correlation between anticipatory and post-meal insulin, without removing non-responders, shows similar results (n=17, R=0.62, p=0.0009). (**H**) Re-analysis of the effect of InsCtx optogenetic inhibition on post-meal insulin levels, without removing non-responders, shows a similar result (n_tdTomato_=9, n_ChrimsonR_=8, p_’Light control’_=0.012, p_’opsin control’_=0.002). (**I**) Blood glucose levels at baseline and during the anticipatory phase, on the day of control illumination (tdtomato, Light ON), no light inhibition (ChrimsonR, control) and on the day with light on during the anticipatory phase (InsCtx inhibition, ChrimsonR, Light ON). There was a significant increase in blood glucose during the ‘light ON’ control day (n=9, p=0.008) and during the InsCtx inhibition day (n= 8, p = 0.002). Note that this variability in increased blood glucose in different days is likely due to the prolonged waiting period in our optogenetic experiments and cannot explain the decrease in anticipatory insulin upon optogenetic inhibition of InsCtx (see ‘L’ and Supp. Fig. 4B,C). **(J-K)** Mice with no anticipatory phase insulin release during control experiments of the optogenetic protocols (**J**) still had significant increase in blood glucose (**K**, n=15, p=0.0003), likely related to stress-induced hyperglycemia in optogenetic experiments. (**L**) No correlation between blood glucose levels and insulin levels during the anticipatory phase (n_tdTomato_=9, n_ChrimsonR_=8, R = 0.3, p = 0.14), indicating that the increase in blood glucose levels in the optogenetic inhibition day is not the reason for the lack of anticipatory insulin release. As mentioned in ‘I’, there should have been a negative correlation in that case. (**M**) Whole blood glucose curve. Glucose levels at 30 minutes post-meal onset show significantly less clearance (n=8, p = 0.045), suggesting glucose homeostasis impairment. Data are presented as individual measurements (points) and mean (vertical bars) ± SEM. Gray dashed lines represent the same mouse. P values were calculated by Student’s paired two sample t-test (for **E,F,G,I,J,K**), independent Student’s t-test (for **H**), Wilcoxon (for **E,H,I**), Mann Whitney’s U test (for **H**), Benjamini-Hochberg correction (**H**), Spearman’s rank test (for **G,L**). *p<0.05, **p <0.01, ***p<0.001, *****p <0.00001.

**Supp. Fig. 4:**
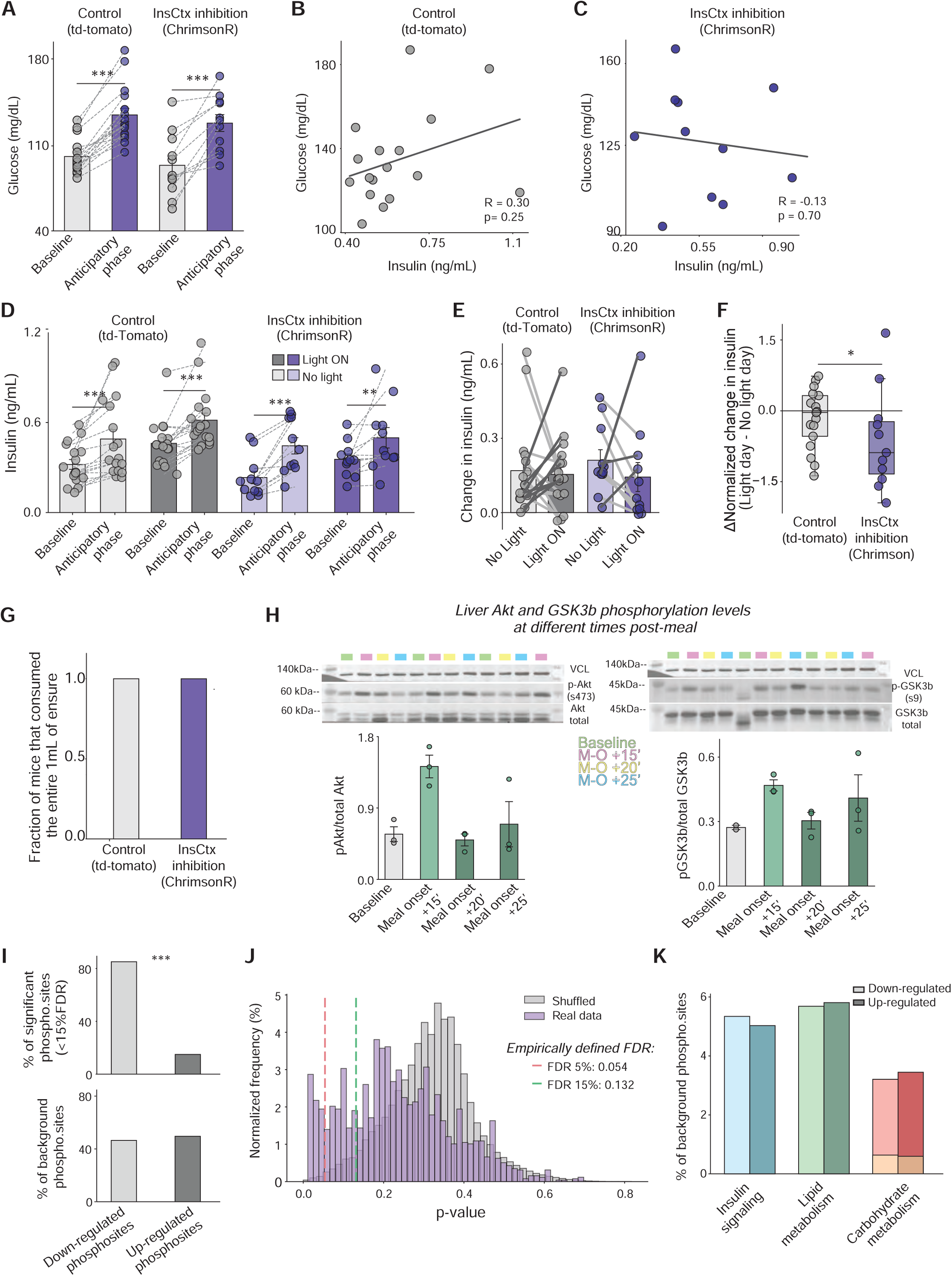
Further control experiments and analyses for across-subjects optogenetic inhibition experiments. **(A)** Blood glucose levels at baseline and during the anticipatory phase for both experimental groups in the ‘light ON’ experimental day. There was a significant increase in blood glucose for both groups in the anticipatory phase (n_tdTomato_ = 17, n_ChrimsonR_ = 11, p-value_tdTomato_ = 0.0003, p-value_ChrimsonR_ = 0.0004). **(B-C)** No correlation between blood glucose and insulin levels during the anticipatory phase for the control cohort (‘B’; n=17, R = 0.3, p = 0.25) and inhibition cohort (‘C’; n=11, R = −0.13, p = 0.7), indicating that the increase in blood glucose in the optogenetic inhibition day does not explain anticipatory insulin release. (**D**) Raw insulin values for all experimental groups in the across-subjects optogenetic experiment (for harvesting tissue). Control group (dTomato, n=17) showed the expected increase in both experimental days (p_No light_= 0.00003 and p_Light on_=0.0002). The experimental group (ChrimsonR, n=11) showed the expected increase during the ‘No light’ day but an attenuated response during the ‘Light ON’ day (p_No light_= 0.0007 and p_Light on_=0.002). (**E**) Change in insulin levels in the across-subjects experiment between the optogenetic inhibition day and the control day for the control group (expressing tdTomato), and inhibition group (expressing ChrimsonR). There was a trend towards attenuation of anticipatory insulin release during the optogenetic inhibition day only in the ChrimsonR mice cohort (8/11 mice, p=0.12, not significant due to one subject with a very strong opposite effect, see ‘F’). (**F**) Normalization of the data from ‘E’ to each mouse’s baseline levels shows a significant decrease in anticipatory insulin upon InsCtx inhibition. To correct for the high variability observed in the data, we normalized it to baseline [(Anticipatory - Baseline)/Baseline]. This showed an effect of InsCtx inhibition in the ChrimsonR cohort but not in the tdTomato cohort (n_tdTomato_ = 17, n_ChrimsonR_ = 11, p-value = 0.037). (**G**) All mice consumed the total amount of Ensure given (1 mL) in all experimental days (n_tdTomato_ = 17, n_ChrimsonR_ = 11). (**H**) Western blot analysis of phosphorylation levels of liver Akt and GSK3b on specific phosphosites (s473 and s9, respectively) at different timepoints (Baseline, and 15, 20 and 25 minutes after start of consumption of 1 mL Ensure, lanes for each category are green (B: baseline), pink (M-O +15’: meal onset +15’), yellow (M-O +20’: meal onset +20’) and light blue (M-O +25’: meal onset +25’)). The highest level of phosphorylation for both phosphosites was at 15 minutes post meal onset. We therefore chose this timepoint for tissue collection in following experiments. (**I**) *Top:* Percentage of significant phosphosites that are down-regulated or up-regulated out of all significant phosphosites (FDR<15%). (Fischer’s exact test. p = 3.57e-50 vs. background. *Bottom:* Percentage of phosphosites that are down-regulated or up-regulated in “background phosphosites”, defined as datapoints with a log2(foldchange) between −0.5 and 0.5. (**J**) Histogram of adjusted p-values for log2(fold change) of shuffled data with 1000 permutations randomly divided into two groups of n=11 and n=17 (gray) and adjusted p-values of original data (purple). Normalized frequency= counts in bin/total counts * 100. Red line: p-value=0.0541 with 5% FDR, and green line: p-value 0.13 with 15% FDR. (**K**) Percentage of phosphosites related to different aspects of metabolism in the background phosphosites (datapoints with a log2(fold change) between −0.5 and 0.5). Insulin signaling (blue), Lipid metabolism (green), Carbohydrate metabolism (red) and Gluconeogenesis (yellow), light shade indicates down-regulation and darker shade up-regulation. There was no significant bias towards either up-regulated or down-regulated phosphorylation. Data are presented as individual measurements (points) and mean (vertical bars) ± SEM. Gray dashed lines represent the same mouse. P values were calculated by Student’s paired two sample t-test (for **A**), Spearman’s rank test (for **B**), Pearson R (for **C**), Wilcoxon (**E**), Mann-Whitney (**F**), Fischer’s exact test (for **I,K**). *= p<0.05, ***** = p <0.00001.

## References

1. Pavlov, I. P. The Work of the Digestive Glands. (C. Griffin, London, 1910).

2. Smith, G. P. Pavlov, and integrative physiology. American Journal of Physiology-Regulatory, Integrative and Comparative Physiology 279, R743–R755 (2000).

3. Cannon, W. B. The Wisdom of the Body, 2nd Ed. (Norton & Co., Oxford, England, 1939).

4. Ramsay, D. S. & Woods, S. C. Physiological Regulation: How It Really Works. Cell Metabolism 24, 361–364 (2016).

5. Barrett, L. F. & Simmons, W. K. Interoceptive predictions in the brain. Nat Rev Neurosci 16, 419–429 (2015).

6. Paulus, M. P., Feinstein, J. S. & Khalsa, S. S. An Active Inference Approach to Interoceptive Psychopathology. Annu. Rev. Clin. Psychol. 15, 97–122 (2019).

7. Pezzulo, G., Rigoli, F. & Friston, K. J. Hierarchical Active Inference: A Theory of Motivated Control. Trends in Cognitive Sciences 22, 294–306 (2018).

8. Quadt, L., Critchley, H. D. & Garfinkel, S. N. The neurobiology of interoception in health and disease. Annals of the New York Academy of Sciences 1428, 112–128 (2018).

9. Gogolla, N. The insular cortex. Current Biology 27, R580–R586 (2017).

10. Saper, C. B. The Central Autonomic Nervous System: Conscious Visceral Perception and Autonomic Pattern Generation. Annual Review of Neuroscience 25, 433–469 (2002).

11. Livneh, Y. & Andermann, M. L. Cellular activity in insular cortex across seconds to hours: Sensations and predictions of bodily states. Neuron 109, 3576–3593 (2021).

12. Livneh, Y. et al. Estimation of Current and Future Physiological States in Insular Cortex. Neuron 105, 1094–1111.e10 (2020).

13. Meier, L. et al. Thirst-Dependent Activity of the Insular Cortex Reflects its Emotion-Related Subdivision: A Cerebral Blood Flow Study. Neuroscience 383, 170–177 (2018).

14. Gardner, M. P. H. & Fontanini, A. Encoding and Tracking of Outcome-Specific Expectancy in the Gustatory Cortex of Alert Rats. J. Neurosci. 34, 13000–13017 (2014).

15. Livneh, Y. et al. Homeostatic circuits selectively gate food cue responses in insular cortex. Nature 546, 611–616 (2017).

16. Samuelsen, C. L., Gardner, M. P. H. & Fontanini, A. Effects of cue-triggered expectation on cortical processing of taste. Neuron 74, 410–422 (2012).

17. Gehrlach, D. A. et al. Aversive state processing in the posterior insular cortex. Nat Neurosci 22, 1424–1437 (2019).

18. Vincis, R., Chen, K., Czarnecki, L., Chen, J. & Fontanini, A. Dynamic Representation of Taste-Related Decisions in the Gustatory Insular Cortex of Mice. Current Biology 30, 1834–1844.e5 (2020).

19. de Araujo, I. E. & Simon, S. A. The gustatory cortex and multisensory integration. Int J Obes 33, S34–S43 (2009).

20. Jones, L. M., Fontanini, A. & Katz, D. B. Gustatory processing: a dynamic systems approach. Current Opinion in Neurobiology 16, 420–428 (2006).

21. Nicolas, C. et al. Linking emotional valence and anxiety in a mouse insula-amygdala circuit. Nat Commun 14, 5073 (2023).

22. Schiff, H. C. et al. Experience-dependent plasticity of gustatory insular cortex circuits and taste preferences. Science Advances 9, eade6561 (2023).

23. Hashimoto, K. & Spector, A. C. Extensive Lesions in the Gustatory Cortex in the Rat Do Not Disrupt the Retention of a Presurgically Conditioned Taste Aversion and Do Not Impair Unconditioned Concentration-Dependent Licking of Sucrose and Quinine. Chemical Senses 39, 57–71 (2014).

24. Kusumoto-Yoshida, I., Liu, H., Chen, B. T., Fontanini, A. & Bonci, A. Central role for the insular cortex in mediating conditioned responses to anticipatory cues. Proceedings of the National Academy of Sciences 112, 1190–1195 (2015).

25. Stern, S. A. et al. Top-down control of conditioned overconsumption is mediated by insular cortex Nos1 neurons. Cell Metabolism 33, 1418–1432.e6 (2021).

26. Contreras, M., Ceric, F. & Torrealba, F. Inactivation of the interoceptive insula disrupts drug craving and malaise induced by lithium. Science 318, 655–658 (2007).

27. Schier, L. A., Hashimoto, K., Bales, M. B., Blonde, G. D. & Spector, A. C. High-resolution lesion-mapping strategy links a hot spot in rat insular cortex with impaired expression of taste aversion learning. Proceedings of the National Academy of Sciences 111, 1162–1167 (2014).

28. Yiannakas, A. & Rosenblum, K. The Insula and Taste Learning. Front Mol Neurosci 10, 335 (2017).

29. Kayyal, H. et al. Activity of Insula to Basolateral Amygdala Projecting Neurons is Necessary and Sufficient for Taste Valence Representation. J. Neurosci. 39, 9369–9382 (2019).

30. Peng, Y. et al. Sweet and bitter taste in the brain of awake behaving animals. Nature 527, 512–515 (2015).

31. Petzschner, F. H., Garfinkel, S. N., Paulus, M. P., Koch, C. & Khalsa, S. S. Computational Models of Interoception and Body Regulation. Trends in Neurosciences 44, 63–76 (2021).

32. Owens, A. P., Allen, M., Ondobaka, S. & Friston, K. J. Interoceptive inference: From computational neuroscience to clinic. Neuroscience & Biobehavioral Reviews 90, 174–183 (2018).

33. Buijs, R. M., Chun, S. J., Niijima, A., Romijn, H. J. & Nagai, K. Parasympathetic and sympathetic control of the pancreas: A role for the suprachiasmatic nucleus and other hypothalamic centers that are involved in the regulation of food intake. Journal of Comparative Neurology 431, 405–423 (2001).

34. Carpenter, R. H. S. Homeostasis: a plea for a unified approach. Advances in Physiology Education 28, 180–187 (2004).

35. Walker, S. J., Goldschmidt, D. & Ribeiro, C. Craving for the future: the brain as a nutritional prediction system. Current Opinion in Insect Science 23, 96–103 (2017).

36. Power, M. L. & Schulkin, J. Anticipatory physiological regulation in feeding biology: Cephalic phase responses. Appetite 50, 194–206 (2008).

37. Feldman, M. & Richardson, C. T. Role of thought, sight, smell, and taste of food in the cephalic phase of gastric acid secretion in humans. Gastroenterology 90, 428–433 (1986).

38. Stricker, E. M. & Hoffmann, M. L. Presystemic signals in the control of thirst, salt appetite, and vasopressin secretion. Physiology & Behavior 91, 404–412 (2007).

39. Langhans, W., Watts, A. G. & Spector, A. C. The elusive cephalic phase insulin response: triggers, mechanisms, and functions. Physiological Reviews 103, 1423–1485 (2023).

40. Pullicin, A. J., Glendinning, J. I. & Lim, J. Cephalic phase insulin release: A review of its mechanistic basis and variability in humans. Physiology & Behavior 239, 113514 (2021).

41. Ahrén, B. & Holst, J. J. The cephalic insulin response to meal ingestion in humans is dependent on both cholinergic and noncholinergic mechanisms and is important for postprandial glycemia. Diabetes 50, 1030–1038 (2001).

42. Teff, K. L. & Engelman, K. Oral sensory stimulation improves glucose tolerance in humans: effects on insulin, C-peptide, and glucagon. *American Journal of Physiology-Regulatory*, Integrative and Comparative Physiology 270, R1371–R1379 (1996).

43. Powley, T. L. The ventromedial hypothalamic syndrome, satiety, and a cephalic phase hypothesis. Psychological Review 84, 89–126 (1977).

44. Wiedemann, S. J. et al. The cephalic phase of insulin release is modulated by IL-1β. Cell Metab 34, 991–1003.e6 (2022).

45. Glendinning, J. I., Lubitz, G. S. & Shelling, S. Taste of glucose elicits cephalic-phase insulin release in mice. Physiology & Behavior 192, 200–205 (2018).

46. Teff, K. L., Mattes, R. D., Engelman, K. & Mattern, J. Cephalic-phase insulin in obese and normal-weight men: Relation to postprandial insulin. Metabolism 42, 1600–1608 (1993).

47. Kalsbeek, A., la Fleur, S. & Fliers, E. Circadian control of glucose metabolism. Molecular Metabolism 3, 372–383 (2014).

48. Oliveira-Maia, A. J. et al. The Insular Cortex Controls Food Preferences Independently of Taste Receptor Signaling. Frontiers in Systems Neuroscience 6, 5 (2012).

49. Araujo, I. E. de, Schatzker, M. & Small, D. M. Rethinking Food Reward. Annual Review of Psychology 71, 139–164 (2020).

50. Glendinning, J. I., Drimmer, Z. & Isber, R. Individual differences in cephalic-phase insulin response are stable over time and predict glucose tolerance in mice. Physiology & Behavior 276, 114476 (2024).

51. Nisi, A. V. et al. Sugar type and route of delivery influence insulin and glucose-dependent insulinotropic polypeptide responses in rats. American Journal of Physiology-Endocrinology and Metabolism 329, E210–E225 (2025).

52. Jung, A.-H. et al. A Subregion of Insular Cortex Is Required for Rapid Taste-Visceral Integration and Consequent Conditioned Taste Aversion and Avoidance Expression in Rats. eNeuro 9, (2022).

53. Talpir, I. & Livneh, Y. Stereotyped goal-directed manifold dynamics in the insular cortex. Cell Reports 43, 114027 (2024).

54. Allen, W. E. et al. Thirst regulates motivated behavior through modulation of brainwide neural population dynamics. Science 364, 253 (2019).

55. Gehrlach, D. A. et al. A whole-brain connectivity map of mouse insular cortex. eLife 9, e55585 (2020).

56. Li, N. et al. Spatiotemporal constraints on optogenetic inactivation in cortical circuits. eLife 8, e48622 (2019).

57. Klapoetke, N. C. et al. Independent optical excitation of distinct neural populations. Nat Methods 11, 338–346 (2014).

58. Vestergaard, M., Carta, M., Güney, G. & Poulet, J. F. A. The cellular coding of temperature in the mammalian cortex. Nature 614, 725–731 (2023).

59. Ayala, J. E. et al. Standard operating procedures for describing and performing metabolic tests of glucose homeostasis in mice. Disease Models & Mechanisms 3, 525–534 (2010).

60. Brandt, C. et al. Food Perception Primes Hepatic ER Homeostasis via Melanocortin-Dependent Control of mTOR Activation. Cell 175, 1321–1335.e20 (2018).

61. Mechanisms of Insulin Action and Insulin Resistance | Physiological Reviews. https://journals.physiology.org/doi/full/10.1152/physrev.00063.2017.

62. Hammerschmidt, P. et al. CerS6-Derived Sphingolipids Interact with Mff and Promote Mitochondrial Fragmentation in Obesity. Cell 177, 1536–1552.e23 (2019).

63. Henschke, S. et al. Food perception promotes phosphorylation of MFFS131 and mitochondrial fragmentation in liver. Science 384, 438–446 (2024).

64. Prescott, S. L. & Liberles, S. D. Internal senses of the vagus nerve. Neuron 110, 579–599 (2022).

65. Sammons, M. et al. Brain-body physiology: Local, reflex, and central communication. Cell 187, 5877–5890 (2024).

66. Zimmerman, C. A. & Knight, Z. A. Layers of signals that regulate appetite. Curr Opin Neurobiol 64, 79–88 (2020).

67. Augustine, V., Lee, S. & Oka, Y. Neural Control and Modulation of Thirst, Sodium Appetite, and Hunger. Cell 180, 25–32 (2020).

68. Andermann, M. L. & Lowell, B. B. Toward a Wiring Diagram Understanding of Appetite Control. Neuron 95, 757–778 (2017).

69. Brüning, J. C. & Fenselau, H. Integrative neurocircuits that control metabolism and food intake. Science 381, eabl7398 (2023).

70. Berntson, G. G. & Khalsa, S. S. Neural Circuits of Interoception. Trends Neurosci 44, 17–28 (2021).

71. Rossi, M. A. & Stuber, G. D. Overlapping Brain Circuits for Homeostatic and Hedonic Feeding. Cell Metabolism 27, 42–56 (2018).

72. Flynn, F. W., Berridge, K. C. & Grill, H. J. Pre- and postabsorptive insulin secretion in chronic decerebrate rats. *American Journal of Physiology-Regulatory*, Integrative and Comparative Physiology 250, R539–R548 (1986).

73. Zhao, Z. et al. Cannabinoids regulate an insula circuit controlling water intake. Current Biology 34, 1918–1929.e5 (2024).

74. Koren, T. et al. Insular cortex neurons encode and retrieve specific immune responses. Cell 184, 5902–5915.e17 (2021).

75. Kayyal, H. et al. Retrieval of conditioned immune response in male mice is mediated by an anterior–posterior insula circuit. Nat Neurosci 28, 589–601 (2025).

76. Tsuneki, H. et al. Food odor perception promotes systemic lipid utilization. Nat Metab 4, 1514–1531 (2022).

77. Shipley, M. T. Insular cortex projection to the nucleus of the solitary tract and brainstem visceromotor regions in the mouse. Brain Research Bulletin 8, 139–148 (1982).

78. Clemmensen, C. et al. Gut-Brain Cross-Talk in Metabolic Control. Cell 168, 758–774 (2017).

79. Campbell, J. E. & Newgard, C. B. Mechanisms controlling pancreatic islet cell function in insulin secretion. Nature reviews. Molecular cell biology 22, 142 (2021).

80. Sternson, S. M. & Eiselt, A.-K. Three Pillars for the Neural Control of Appetite. Annual Review of Physiology 79, 401–423 (2017).

81. Gizowski, C. & Bourque, C. W. The neural basis of homeostatic and anticipatory thirst. Nat Rev Nephrol 14, 11–25 (2018).

82. Prilutski, Y. & Livneh, Y. Physiological Needs: Sensations and Predictions in the Insular Cortex. Physiology 38, 73–81 (2023).

83. Domingos, A. I. et al. Leptin regulates the reward value of nutrient. Nat Neurosci 14, 1562–1568 (2011).

84. Tan, H.-E. et al. The gut–brain axis mediates sugar preference. Nature 580, 511–516 (2020).

85. Glendinning, J. I. et al. Glucose elicits cephalic-phase insulin release in mice by activating K _ATP_ channels in taste cells. *American Journal of Physiology-Regulatory*, Integrative and Comparative Physiology 312, R597–R610 (2017).

86. Stringer, C., Wang, T., Michaelos, M. & Pachitariu, M. Cellpose: a generalist algorithm for cellular segmentation. Nat Methods 18, 100–106 (2021).

87. Burgess, C. R. et al. Hunger-Dependent Enhancement of Food Cue Responses in Mouse Postrhinal Cortex and Lateral Amygdala. Neuron 91, 1154–1169 (2016).

88. Bonin, V., Histed, M. H., Yurgenson, S. & Reid, R. C. Local diversity and fine-scale organization of receptive fields in mouse visual cortex. J Neurosci 31, 18506–18521 (2011).

89. Mukamel, E. A., Nimmerjahn, A. & Schnitzer, M. J. Automated Analysis of Cellular Signals from Large-Scale Calcium Imaging Data. Neuron 63, 747–760 (2009).

90. Elinger, D., Gabashvili, A. & Levin, Y. Suspension Trapping (S-Trap) Is Compatible with Typical Protein Extraction Buffers and Detergents for Bottom-Up Proteomics. J. Proteome Res. 18, 1441–1445 (2019).

91. Wenger, C. D. & Coon, J. J. A Proteomics Search Algorithm Specifically Designed for High-Resolution Tandem Mass Spectra. J. Proteome Res. 12, 1377–1386 (2013).

92. Li, Q. et al. Global Post-Translational Modification Discovery. J. Proteome Res. 16, 1383–1390 (2017).

